# Elevated human impact on islands increases the introduction and extinction status of native insular reptiles

**DOI:** 10.1101/2022.03.11.483632

**Authors:** Wendy A.M. Jesse, Jacintha Ellers, Jocelyn E. Behm, Gabriel C. Costa, S. Blair Hedges, Matthew R. Helmus

**Affiliations:** Department of Ecological Science, Vrije Universiteit Amsterdam, De Boelelaan 1085 Amsterdam, NL 1081 HV; Department of Biology, Temple University, 1925 N. 12th Street Suite 502, Philadelphia, PA 19122; Department of Biology and Environmental Sciences, Auburn University at Montgomery, Montgomery, AL, 36124, USA

## Abstract

Species ranges are changing in the Anthropocene, the ranges of introduced species are expanding, while extinction-prone species are contracting. Introductions and extinctions are both caused by how species respond to human impacts, but it is unknown why the ranges of some species expand and some contract. Here, we test that this opposite response of human impact is due to introduced and extinction-prone species falling at opposite ends of geographic, evolutionary, or ecological trait continua. We constructed a database of native range maps, traits, phylogenetic relationships, and the introduction and extinction-prone status of squamate reptiles with ranges native to the Western Hemisphere. Across >3,000 snake and lizard species (88% of known native squamates), 142 had been introduced elsewhere and 483 were extinction-prone (i.e., extinct, vulnerable, threatened). To explain variation in status, we first tested if the same human-impacted regions in the Americas contained the native ranges of species of either status. Second, we tested for phylogenetic signal in species status. Finally, we tested the explanatory power of multiple trait continua. The native ranges of introduced and extinction-prone reptiles were clustered in island regions with high human impact vs. mainland regions with lower human impact. Phylogenetic signal was weak for status, but introduced and extinction-prone species were clustered in different clades. All geographic and ecological traits that explained both statuses supported the opposite ends hypothesis. Introduced species had larger, edgier ranges, while extinction-prone species had smaller, simpler ranges. Introduced species were mostly herbivorous/omnivorous, while extinction-prone species were mostly carnivorous. Introduced species produced larger clutches, while extinction-prone species were smaller in body size. In the Anthropocene, the naive ranges of introduced and extinction-prone species are in the same human-impacted regions where trait continua, having opposite effects, determine whether species ranges expand or contract in the continuing face of global change.

## Introduction

In the Anthropocene, humans have altered species ranges and reassembled global biogeographic patterns that arose naturally across eons of evolution (Alroy 2015, Capinha et al. 2015, Ceballos et al. 2017). Range contraction and expansion varies across species due to human impact (Pacifici et al. 2020). However, the ultimate “winners” of the Anthropocene are introduced species, which expand into new geographic regions through human-driven dispersal and establishment (Colautti and MacIsaac 2004). While the “losers” are the extinct and threatened species that have experienced severe native range contraction due to human impacts like overharvesting, pollution, climate change, and land-use change (Vitousek et al. 1997, Sax and Gaines 2008, Böhm et al. 2013, Young et al. 2016). Most introductions and extinctions have occurred since the mid 20^th^ Century, a period considered to be the start of the Anthropocene epoch where multiple indicators of human impact accelerated (IPBES 2019, 2023, Waters and Turner 2022).

The reasons for the opposite responses of introduced and extinction-prone species to human impact are multitiered. Species that get introduced to other regions often have native ranges that overlap with dense human populations increasing likelihood of intentional (e.g., pet trade) or unintentional (e.g., stowaways) human-aided dispersal (Latella et al. 2011, Liu et al. 2014, Su et al. 2016, Perella & Behm 2020). After introduction, species with suitable functional traits tend to be more successful in utilizing novel food sources and endure local climatic regimes (Mahoney et al. 2015, Monaco et al. 2020). In contrast, extinction-prone species generally are ecological specialists, characterized by narrow dietary and climatic niches and poor dispersal abilities due to their geographic restriction to mountain tops, islands, or isolated nature reserves in a matrix of anthropogenic land use (Chichorro et al. 2019, Kotiaho et al. 2005, Böhm et al. 2016). Hence, introduced and extinction-prone species appear to be not only each other’s antipodes in how they respond to human influence but also in many characteristics that could explain their opposite range responses in the Anthropocene. This raises the question if these winners and losers are at opposite ends of geographic, evolutionary, and ecological trait continua (Schmidt et al. 2021, Jeschke and Strayer 2008, Blackburn and Jeschke 2009). Testing this introduced-extinct opposite ends of the same trait continua hypothesis (also generically termed the two-sides-of-the-same-coin hypothesis in the literature) requires data on functional traits, native range characteristics, and geographic and phylogenetic clustering associated with introduced and extinction-prone species that are analyzed to determine the causes of dissimilarity between the two groups.

The studies that have tested the introduced-extinct opposite ends hypothesis show variable results. For instance, extinction-prone and successfully introduced plants (Bradshaw et al. 2008, Pandit et al. 2011, Schmidt et al. 2012), fish (Liu et al. 2017), mammals (Pacifici et al. 2020) and crayfish (Larson and Olden 2010) were on opposite extremes of the same trait axes such as body size, fecundity, longevity, genetic diversity, intraspecific trait variation, and habitat specialization. However, not all traits showed the opposite pattern. Other studies on birds and reptiles have found little evidence for the opposite-ends hypothesis even though different groups of traits characterized either extinction-prone or introduced species (Jeschke and Strayer 2008, Blackburn and Jeschke 2009, Tingley et al. 2016, Marino & Bellard 2023). Furthermore, such comparative studies generally lack an (in-depth) phylogenetic perspective on species introduction and extinction probabilities, which is needed because nonoverlapping phylogenetic clustering of introduced and extinct species is expected under the hypothesis because many traits or characteristics of species exhibit phylogenetic signal (Schmidt et al. 2021). Moderate to strong levels of phylogenetic signal have been found in traits associated with introduction and/or invasion success (Cadotte et al. 2009, Park and Potter 2015), native range size (Pigot et al. 2018), and fecundity (Allen et al. 2017, Yessoufou et al. 2016, Pyšek et al. 2017, Alcaraz et al. 2005, Su et al. 2016). Also, phylogenetic clustering of extinction-prone species has been detected in various taxa (Davies et al. 2011, Loza et al. 2017, Adeoba et al. 2019, Arbetman et al. 2017, Fritz and Purvis 2010, Tonini et al. 2016). Overall, there is compelling evidence that the predisposition to becoming introduced or extinct is phylogenetically clustered and should be considered when testing for the opposite ends hypothesis.

How species respond to human influence is assumed to be related to native geographic range characteristics associated with range expansions and contractions. Island-living may be an important characteristic determining species status. Islands have high human population densities and are subjected to disproportionate levels of human impact (Kier et al. 2009), thus insular species might be more likely to be introduced or go extinct than mainland species. Islands are hotspots of species loss (Myers et al. 2000, Mittermeier et al. 2011) and provide unique environments that select for functional traits that naturally determine major range expansions or contractions. Insular species have evolved through processes of oversea colonization, followed by often rapid adaptive diversification (Cowie and Holland 2006, Hedges 2006). Thus, insular species are selected to be efficient dispersers and quick adapters to available niche space, which could favor survival of human-vectored dispersal events (Poe et al. 2011). Indeed, post-introduction ecological niche shifts can occur for introduced species that originate from oceanic islands (Liu et al. 2014, Stroud 2021). However, island adaptive radiations may also cause island biota to be highly specialized, dispersal-limited, and extremely suited to exploit a narrow ecological niche within island environments (Losos 2009, Mahler et al. 2010, 2013). This level of specialization leaves island endemics sensitive to human-impact and thus prone to extinction (Jantz et al. 2015).

In this study, we used a phylogenetic comparative approach to identify dissimilarities in introduced and extinction-prone species characteristics. We built a dataset of 3111 squamate reptiles, which is 84% of all known lizards and snakes native to the Western Hemisphere (Fig. 1). We identified 142 species that had been introduced to at least one location somewhere in the world and 483 species threatened with extinction according to the International Union for the Conservation of Nature (IUCN 2021, Cox et al. 2022). Reptiles of the Americas are particularly suitable to test opposite responses to human influence, as reptiles are species-rich in the Western Hemisphere and are greatly impacted by human activities (Young et al. 2016, Jesse et al. 2018, Gleditsch et al. 2023). We asked: 1) Are islands of the Western Hemisphere disproportionate sources of introduced and sinks of extinction-prone species? 2) Can we detect phylogenetic signal among introduced and extinction-prone species? 3) For which geographic, evolutionary, and ecological trait continua are introduced and extinction-prone species positioned at opposite ends?

**Figure 1.**
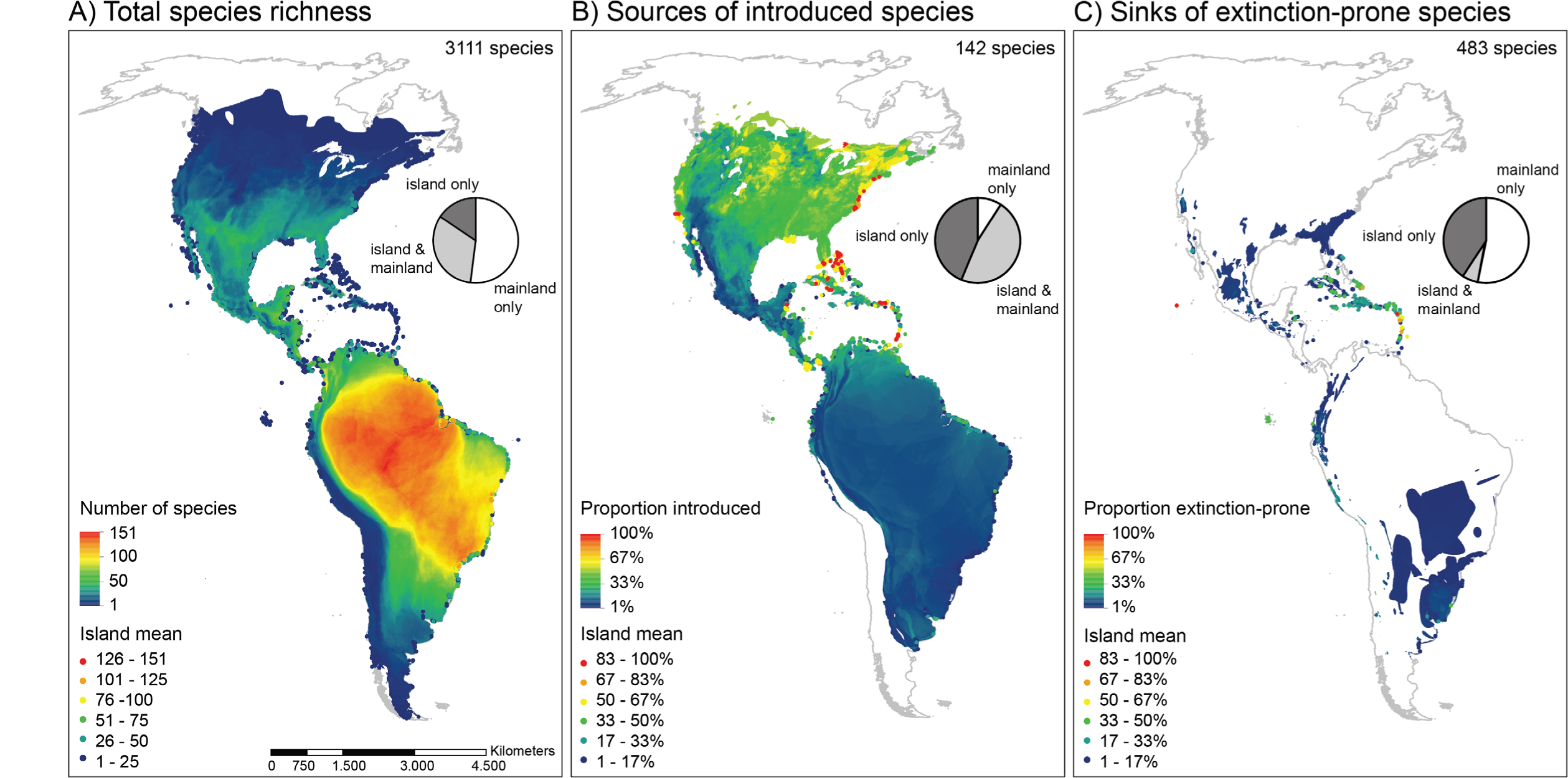
The geographic clustering of the native ranges of introduced and extinction-prone American reptile species indicates islands as sources and sinks of Anthropocene biodiversity. A) The total of overlapping native species ranges per 1km^2^ grid cell provides a map of Western Hemisphere reptile species richness. The proportions of introduced (B) and extinction-prone (C) richness in relation to total species richness were calculated for all 1 km^2^ grid cells inhabited by >1 species. Points in all maps represent mean values per island and follow the same color scheme as the gridded layers. Pie charts represent the geographic context of the species ranges depicted in the respective maps, showing the proportions of species that are restricted to the continental mainland of North and South America (white), partially insular species that inhabit island and mainland areas (light grey), and species that are restricted to oceanic islands (dark grey). Percentages within pie charts are (clockwise, i.e., *mainland only, island and mainland, island only*) (A) 52%, 32%, 16%, (B) 9%, 47%, 44%, (C) 53%, 6%, 41%. All maps were projected as Eckert IV spherical world projection in ESRI ArcMap 10.6.1.

## Methods

### Data compilation

#### Squamate phylogeny

We updated a global, smoothed, and interpolated phylogeny of squamates from the TimeTree of life project (Kumar et al. 2017), which contained 9378 worldwide species previously built in Marin et al. (2018) and Rapacciuolo et al. (2019). Species synonyms were identified and cleaned using the Reptile Database (Uetz, P., Freed, P. & Hošek 2012) to match up with IUCN native range polygons (see “Geographic data” below). Species not in the phylogeny and not in the spatial dataset were not analyzed. Specifically, the IUCN listed 3519 Western Hemisphere species of which 3463 had native range distributions available. Intersecting the phylogeny with the range polygons, resulted in a phylogeny of 3111 of species (Fig. 2, 88% of known Western Hemisphere species). We analyzed this subset of species for phylogenetic signal in introduced and extinction-prone status. Of these 3111 species, 2936 had IUCN native range maps and data on traits. We analyzed this subset of species in multivariable phylogenetic generalized linear models (PGLMM) to test if introduced and extinction-prone species fall at opposite ends of the same trait continua.

**Figure 2.**
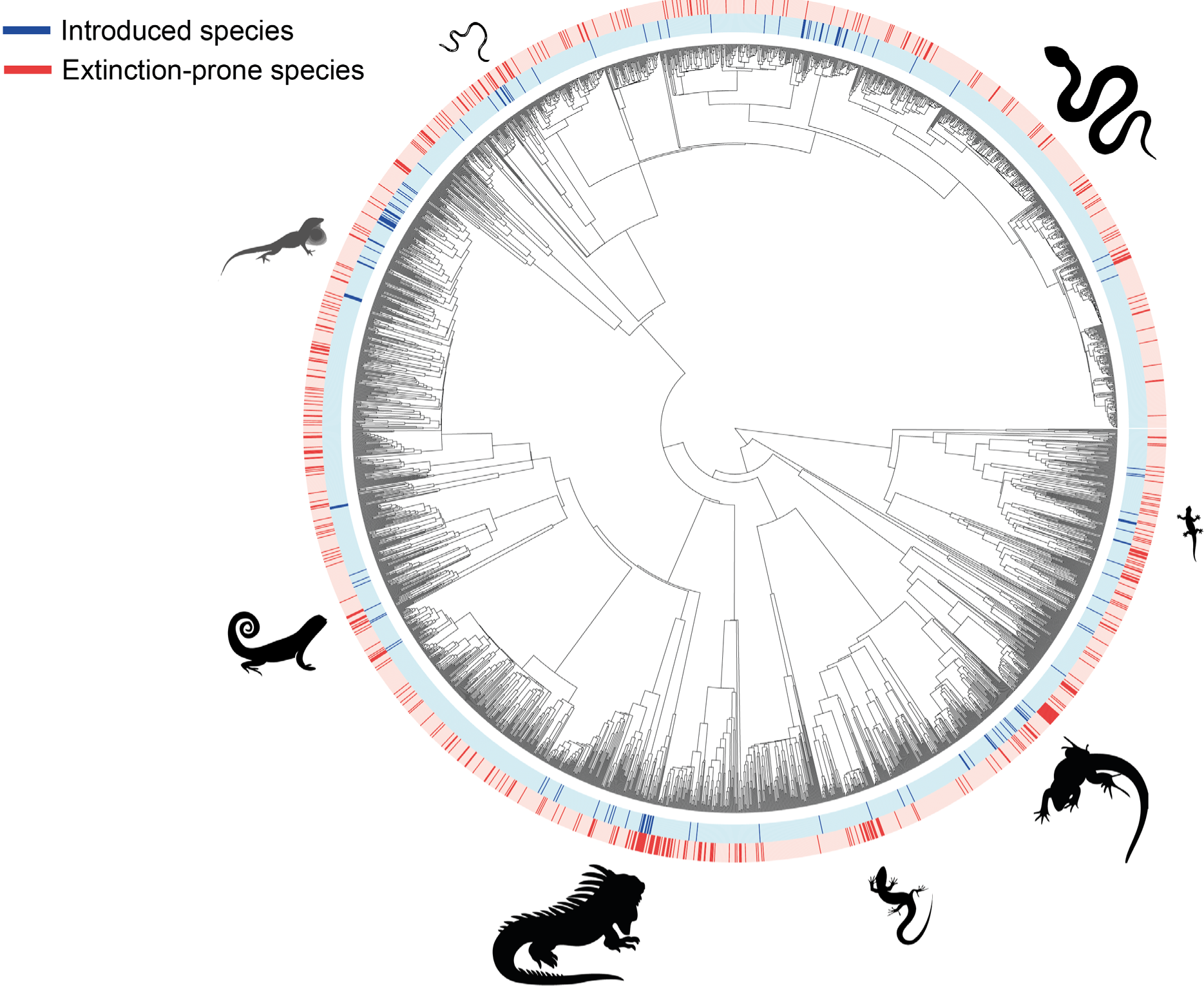
The phylogenetic clustering of introduced and extinction-prone status of American reptiles indicates they are not a random subset of all the reptiles that have evolved in the Western Hemisphere. Introduced species (142) and extinction-prone species (483) are clustered across the Western Hemisphere squamate tree-of-life (3111 species total). Species with an introduced status were those with at least one population established outside of their native range in the Americas. Species with an extinction-prone status were categorized by the IUCN as “vulnerable”, “endangered”, “critically endangered”, “extinct in the wild”, and “extinct.” Silhouettes are of genera with clusters of introduced and/or extinction-prone species (clockwise: *Boa, Sphaerodactylus, Ameiva, Scincus, Iguana, Cnemidophorus, Anolis, Typhlops*). A metric of phylogenetic signal that indicates clustering (D = 0) or a random distribution (D = 1) for introduced was D = 0.63 (P < 0.001 compared to both distributions) and for extinction-prone was D = 0.71 (P < 0.001).

#### Species introduction and extinction-prone status

The response variables in our analyses were based on species statuses of squamates across the Western Hemisphere. We constructed a binomial variable to indicate whether a species was introduced anywhere on earth, and a response variable that indicated if a species was extinction-prone or not. To determine introduction status, we cross-referenced 3111 species from the phylogeny for which we had spatial native range data (see “Geographic data” below) with published invasive species databases, such as the Global Register of Introduced and Invasive Species, the Global Invasive Species Database, the Invasive Species Compendium, and others (e.g., Powell et al. 2011, Mahoney et al. 2015, Tingley et al. 2016, Helmus et al. 2017, Behm et al. 2022; see Table S1 for the full list of references). For all species that did not match with these databases, we performed a Google query using the search term “*Species name* AND introduced OR exotic OR invasive OR alien OR nonnative OR non-indigenous” to look for species-specific literature. We excluded species for which introduction records were not scientifically published, uncertain, or debated. Extinction risk was taken from IUCN records (accessed August 2018). Species that were “vulnerable”, “endangered”, “critically endangered”, “extinct in the wild”, and “extinct” were all categorized as extinction-prone. Of the 3111 species, 349 of category “data deficient” and 494 of category “not assessed” were excluded from the analyses (see “Analyses” below).

#### Geographic data

Distributional range maps for 3463 reptile species were compiled from the IUCN and Caribherp databases (Hedges 2021, IUCN 2021) and edited to only include polygon shapes of native ranges of squamates in the Western Hemisphere that also appeared in the phylogenetic tree. The resultant dataset included 3111 species. This difference in the number of species between datasets was caused by the mismatch between phylogeny and geographic ranges and removal of duplicate ranges, taxonomic synonyms, exotic ranges, and non-squamate species. We used these range polygons to calculate covariates (italicized subheadings below) for our statistical analyses.

##### Range size and shape

For each species, we calculated native range size and shape. Range size was estimated as range area in km^2^. Range shape was estimated as the inverse of the normalized perimeter of the range, which is calculated as range perimeter divided by the perimeter of a circle with the same area as the range (Patton 1975). While there are many metrics of range shape, most use some equation that includes a measure of range perimeter and area often expressed as a ratio (Krummel et al. 1987, Kupfer 2012). We chose normalized range perimeter (i.e., also termed edge diversity index, perimeter-area ratio shape index) because it is an intuitive metric indicative of range edginess versus compactness that is widely used and statistically independent of range size (Frazier and Kedron 2017). Values closer to one indicate ranges with many edges such as those that include multiple geographic boundaries. Values closer to zero are more compact and circular. Range size and shape were estimated with the “areaPolygon” (m^2^) and “perimeter” (m) functions from the geosphere R package (Hijmans 2019). We expected that a species is more likely to be introduced if it has a larger range with more edges. Larger ranges means there is more area for the species to hitchhike along transportation networks. Similarly, more edges means the species is distributed across multiple islands, found near coastlines, in river valleys, and more people live near water (Small & Nicholls 2003). In contrast, we expected extinction-prone species to have small and simpler ranges, confined to a single or few populations.

##### Species insularity

For each species we calculated its level of insularity as the portion of a species range located on oceanic islands. We intersected species range polygons with a map of the continental mainland of North and South America (GADM 2018), enabling us to identify the mainland continental ranges for all species. Based on the area difference between the total species range and the continental range, we were able to infer the insular fraction of a species range. An insularity value of 1 indicates exclusive island living (i.e., island endemic) and values of 0 indicated that species only occur on the continental mainland. We expected both introduced and extinction-prone species to be native to islands because islands are highly populated and impacted by humans compared to the mainland.

#### Evolutionary data

##### Genus age

Genus age was determined by calculating the branching-time from the most recent common ancestor of the genus (i.e., the stem age) with the ‘AssessMonophyly’ and ‘branching.time’ functions (‘ape’ R package). Of 280 Western Hemisphere genera, 212 were monophyletic, 65 were monotypic (i.e., a genus containing one species) and three were paraphyletic (*Epicrates*, *Leposoma*, *Homonota*). The intruder or outlier clades in paraphyletic genera were all monophyletic or monotypic, thus the distribution of genus ages was unaffected. We expected genus age to positively associated with introduced status, and negatively associated with extinction-prone status because of its observed relationship to niche volumes and range expansions (Davies et al. 2011, Title & Burns, 2015).

##### Evolutionary range expansion rate

We summed the species range areas for all the species within the same genus to represent the cumulative range area that a genus has spread into since the time of divergence from the most recent common ancestor. Cumulative range area was divided by the focal genus age to calculate the average evolutionary range expansion rate of geographic spread for all species within a genus. We expected this metric to be negatively associated to extinction-prone status based on previous relationships found for reptiles (Title & Burns 2015), and positively associated with introduced status because squamate lineages that were successful colonizers of ancient Caribbean islands also tend to have more introduced species (Poe et al., 2011).

#### Ecological data

##### Functional traits

We obtained data on functional traits hypothesized to be related to opposite distributional responses to human influence (Jeschke and Strayer 2008, Blackburn and Jeschke 2009, Tingley et al. 2016). Trait data were from Rapacciuolo et al. (2019). Specifically, we expected clutch size, viviparity, omnivory, and body size to be positively (negatively) associated with introduction (extinction-prone) status. Natural log-transformed values of clutch size and maximum body size, as well as two categorical variables for diet (herbivorous, carnivorous, and omnivorous) and reproductive mode (binomial: oviparous and (partially) viviparous), were included as independent variables in PGLMs.

##### Seasonality and elevation

We used a Principal Component Analysis (PCA) to extract composite variables that best explain the climatic variation experienced by species. First, we calculated the median of all 37 climatic variables in WorldClim and ENVIREM raster layers (30 arcsecond resolution, ca. 1km^2^) per species range (Fick and Hijmans 2017, Title and Bemmels 2018). Subsequently, we included all range medians in a scaled PCA in R, of which the first two principal components, together explaining 70% of median environmental variation, were selected. The first axis (PC1) aligned with climatic seasonality variables such as annual temperature range, diurnal temperature range, and seasonal variation in potential evapotranspiration, and the second axis (PC2) aligned with two elevation-derived variables from the ENVIREM dataset: terrain roughness and topographic wetness (indicative for the catchment of water in a watershed) (Title and Bemmels 2018; See Fig. S1). We expected extinction probability to be higher for species associated to isolated mountain ranges (e.g. Guirguis et al., 2023) and introduced species to be associated with seasonal climates with broad environmental tolerances (e.g. Tingley et al. 2016).

##### Climatic niche differentiation

We assessed the climatic variability in species ranges as indication of a species’ adaptive capacity to various climatic regimes, expecting introduced species to have a high capacity and extinction-prone species to have a low capacity. We calculated the standard deviations of the 37 WorldClim and ENVIREM climatic variables per species range and included these in a scaled PCA. PC1 of this PCA (explaining 63% of all variation) was positively correlated to variability for all climatic variables (Fig. S2). Therefore, the PC1 score per species was taken as a value for climatic variability per species range, which was positively correlated with log-transformed range area (r = 0.65, t_3054_ = 47.45, *P* < 0.001), that is, species with large ranges experience high levels of climatic variation. We used the residuals of this linear relationship to reflect climatic niche differentiation relative to a species range size.

#### Anthropogenic data

##### Human footprint and number of ports

As a general indicator of human impact within species ranges, we calculated the median human footprint for every species range (30 arcsecond resolution, ca. 1km^2^), which is a compilation of the amount of built environment, population density, agriculture, and several types of terrestrial and waterborne infrastructure (Venter et al. 2016). The median value was calculated across all pixels within each species range. We expected positive associations of human footprint with both introduced and extinction-prone status. We used the number of sea ports as a human impact indicator of propagule pressure to and from native ranges. Firstly, ports can serve as points of species export, positively affecting introduction status (Hulme, 2009). Number of ports in the native range also explains naturalization success of anole lizards (Latella et al. 2011). Secondly, ports can serve as points of entry of exotic species (Schneider et al., 2021), also promoting extinction-prone status, because native species become subjected to negative species interactions form these exotics. We determined the number of ports (incl. harbors, seaports, and major terminals) per native range area (World Port Index; National Geospatial-Intelligence Agency 2017) as a proxy for the level of seaborne trade to and from the species native range. Number of ports and human footprint index were not significantly correlated (r=-0.02, t_2988_=-0.98, *P*=0.33), and therefore both were included in the statistical analyses.

### Analyses

All analyses were performed in R. We first asked if islands are sources of introduced species and sinks of extinction-prone species. We made maps of squamate species richness and proportions of introduced and extinction-prone richness in relation to total species richness using 1 km^2^ grid cells. We then calculated the proportion of species with introduced or extinction-prone status whose ranges were found only on the mainland, mainland and islands, or only islands; and tested if species insularity explained introduced and extinction-prone status with chi-squared tests.

Second, we asked if there was phylogenetic signal among introduced and extinction-prone species indicating clustering in clades of species with similar status. The level of phylogenetic signal among introduced and extinction-prone species was determined with a metric D that tests for signal in binomial data (phylo.d in caper, Orme et al. 2013). We chose this metric because it exhibits statistically robust expectations for large trees when the proportion of species with a status value is low, like for our data (Fritz & Purvis 2010). The observed D value for introduced and extinction-prone species were each compared to two null distributions. First, to test if there was any phylogenetic signal in the data that is different from random and indicative of phylogenetic clustering, a distribution of 1000 random D values was generated under a null model permutation that randomized status values among species irrespective of the phylogeny. Second, to test the strength of clustering, a distribution of 1000 D values were generated from a null model that simulated status according to Brownian evolution across the phylogenetic tree. Observed D values different than the random null indicated clustering, while values that fell between the two distributions indicated weak clustering.

Finally, we tested the introduced-extinct opposite ends hypothesis by asking which geographic, evolutionary, and ecological trait continua explained introduction and extinction-prone status. The trait continua exhibited phylogenetic signal (Table S2), thus we used binomial phylogenetic generalized linear models (PGLMs) with logistic error term and Firth’s penalized likelihood correction (phyloglm in phylolm, Ho & Ane 2014). First, two separate PGLMs with introduction and extinction status as dependent variables were performed with range insularity as the sole independent factor, because of collinearity between insularity and other covariates (Fig. S3). All other covariates had relatively low levels of collinearity (r≤|0.60| and VIF < 3.0). Second, we ran two separate PGLMs with the other covariates not including insularity to test the hypothesis. Prior to all analyses, we scaled the covariates (mean = 0, sd = 1), so model estimates represented effect sizes.

## Results

The native ranges of introduced and extinction-prone squamate species of the Western Hemisphere were often found on islands (Fig. 1). Across all species, 52% had native ranges only on the mainland, primarily Amazonia (Fig. 1A). In contrast, 91% of the introduced species were either island endemics (44%) or had islands in their native range (47%, Fig 1B), making islands key sources of introduced reptiles globally. Similarly, island endemics constituted 41% of the extinction-prone species (Fig. 1C). This clear overrepresentation of island endemics in introduced (χ^2^ = 47.83, df = 1, *P* < 0.001) and extinction-prone reptile groups (χ^2^ = 164.67, df = 1, *P* < 0.001) indicates that islands are simultaneous sources and sinks of biodiversity in the Anthropocene.

Introduced and extinction-prone species were phylogenetically clustered across the phylogeny of Western Hemisphere reptiles (Fig 2). The signal for introduced (D = 0.63) and extinction-prone (D = 0.71) statuses both were D < 1 (*P <* 0.001) indicating that statuses more clustered than expected if statuses were randomly distributed across the phylogeny. However, both were D > 0 (*P <* 0.001). Zero is the D value expected under Brownian motion evolution of statuses; thus, the clade clustering of statuses was weak. Introduced and extinction-prone species came from different clades because the composition of the two groups showed little overlap. Of the 142 introduced and 483 extinction-prone squamates, only 15 species were included in both categories. Of these 15 species, eight are popular species in the pet trade and threatened with extinction in their native range (e.g., *Cyclura* iguanas).

Range insularity was positively related to both introduction (0.39 ± 0.08, z = 4.45, *P* < 0.001) and extinction probabilities (0.54 ± 0.05, z = 10.52, *P* < 0.001; Fig. 3A). Range insularity showed strong collinearity with other variables in our analyses, most prominently with range area (r = −0.39, t = −23.3, *P* < 0.001), diet type (Kruskal-Wallis: χ^2^ = 18.2, df = 2, *P* < 0.001; bias towards omnivory), and human footprint (r = 0.46, t = 29.1, df = 3056, *P* < 0.001) (see Fig. S3 for pairwise correlations). This indicates that island-living can be seen as a trait “syndrome”: a set of correlations among individual traits and species-level characteristics (e.g., Poe et al. 2011). Therefore, in our PGLM analyses range insularity was excluded from the set of trait continua used to test if introduced and extinction-prone species are positioned at opposite ends. After exclusion of range insularity from the models, human footprint was the only other explanatory variable that was positively related to introduction and extinction probability (Fig. 3B). The underlying cause of this is likely to be the significantly higher human footprint in species ranges located on oceanic islands than on the mainland (t = 22.03, df = 653.79, *P* < 0.001; Fig. 3C), as well as in the ranges of introduced and extinction-prone species compared to other Western Hemisphere reptiles (t = 3.76, df = 145.41, *P* < 0.001 and t = 11.94, df = 579.02, *P* < 0.001, respectively; Fig. 3C).

**Figure 3.**
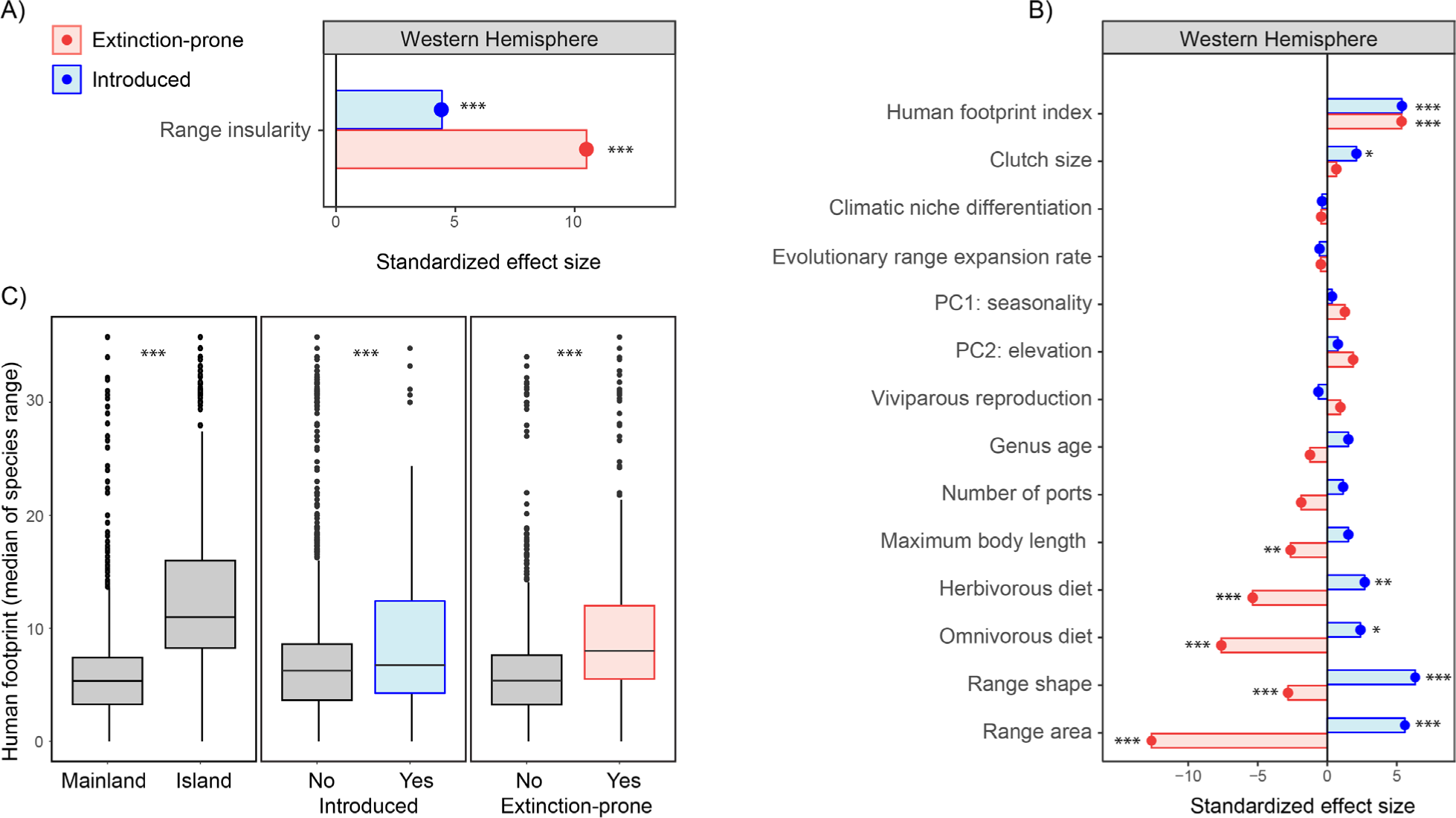
Elevated human impact to islands increases introduction and extinction status of insular lizards and snakes situated at opposite ends of trait continua. A) Across Western Hemisphere squamate reptiles, species with more insular ranges were more likely to be introduced or extinction-prone. B) Species with introduced or extinction-prone status fell on opposite ends of the trait continua that explained both statuses. C) The native ranges of island endemics (Island) are more impacted by humans than continental species (Mainland), and the native ranges of species with introduced or extinction-prone status are also more impacted. In A and C, range insularity was the proportion of a species native range found only on islands and ranged between 1 (island endemics, Island in C) and 0 (continental species, Mainland in C). In A and B, bars are standardized effect sizes (coefficient/SE). In A, separate univariate logistic PGLMMs for introduced or extinction-prone status were fit only to insularity. In B, separate multivariate PGLMMs were fit to a traits based on species geography, evolution, and ecology. In C, the three panels are box plots of the observed data and significance assessed with separate logistic PGLMMs on the data plotted in each panel (*P* < 0.05*, *P* < 0.01**, *P* < 0.001***).

All the trait continua that explained both statuses supported the opposite ends hypothesis (Fig. 3B). Native ranges of introduced species were larger (0.78 ± 0.14, z = 5.58, *P* < 0.001) and more edgy (0.74 ± 0. 12, z = 6.32, *P* < 0.001). Extinction-prone ranges were smaller (−0.86 ± 0.07, z = −12.64, *P* < 0.001) and relatively circular (−0.15 ± 0.05, z = −2.83, *P* = 0.005). Introduced and extinction-prone species had opposing diets. Introduced species were herbivorous (1.27 ± 0.47, z = 2.70, *P* = 0.007) or omnivorous (0.68 ± 0.28, z = 2.39, *P* = 0.02) compared to carnivorous (i.e., carnivory was the contrast in the PGLMs). Conversely, extinction-prone species were carnivorous rather than herbivorous (−2.19 ± 0.41, z = −5.37, *P* < 0.001) or omnivorous (−1.70 ± 0.22, z = −7.62, *P* < 0.001). Seven of the tested trait continua were uninformative, but two others explained a single status (Fig 3B). As expected, introduced species had larger clutches (0.25±0.12, z=-2.10, *P*=0.04), but unexpectedly, extinction-prone species were smaller (−0.11±0.04, z=-2.63, *P*=0.008).

## Discussion

Islands contain the most human-dominated ecosystems on earth, and island living today impacts the status of native lizards and snakes across the Western Hemisphere. Our results suggest that native squamate species on islands have a higher likelihood of being either introduced or extinction-prone. We found that phylogeny predicted status, but the signal was weak. Instead, ecological and geographic trait continua best explained status. While species of either status were insular, trait continua oppositely affected introduction vs extinction risk of species to human impact. Below we discuss each of our major results. The results allow for a better understanding of range dynamics under global change and why highly impacted oceanic islands are simultaneous sources and sinks of biodiversity in the Anthropocene.

### Insularity of introduced and extinction-prone species

Islands have consistently been found to have more established alien species than the mainland (e.g., Li et al. 2023). What we found here was that islands also generate more introduced species than the mainland. In the Western Hemisphere, squamate species with the most insular native ranges were also most likely to be introduced elsewhere (Fig. 1B, 3A). Such a strong relationship has not been documented before to our knowledge, but we did find two relevant studies. First at the global scale, birds that have been introduced and birds that are extinction-prone due to invasive predatory species often have ranges that encompass islands (Marino & Bellard 2023). However, this effect was weak for introduced birds and the authors only studied extinction-prone species impacted by invasives. Second, global spread rates of introduced herpetofauna species are not related to insularity (Liu et al. 2014). However, this study did not compare introduced vs. extinction-prone species and instead looked at how spread rate varied across continents and islands. In sum, insularity seems related to the probability a species is introduced, but not necessarily to how quickly an introduced species spreads.

Many of the extinction-prone squamates native to the Western Hemisphere were island endemics, and the more insular a species native range, the more likely it was to be extinction-prone (Fig. 1C, 3A). Globally, island regions are characterized by simultaneous high levels of endemism and extinction risk (Myers et al. 2000, Mittermeier et al. 2012). They are heavily impacted, and most extinctions have and are occurring on islands (Fernández-Palacios et al. 2021). Islands of the Western Hemisphere, like those in the Caribbean, are highly developed and connected by trade (Gleditsch et al. 2023). Thus, we should expect to see a similar influence of insularity on introduction and extinction-prone status for islands in the Eastern Hemisphere that are also highly impacted. The location of a species’ native range and how heavily it is impacted by humans are key determinants of a species fate in the Anthropocene.

### Introduced and extinction-prone species trait continua

We examined how native range characteristics, functional traits, and human impact influenced status, particularly focusing on range geography, evolutionary history, climate, diet, and life history traits. Human footprint in native ranges significantly raised the chances of a species being introduced or facing extinction, especially for island species (Fig. 3BC). However, we did not detect an association with our other metric of human impact, number of ports in the native range. This is surprising because trade increases propagule pressure of squamates and explains introduced squamates richness (Gleditsch et al. 2023, Mahoney et al. 2015). Port numbers may not well reflect trade pathways for squamates. The pet trade and live-plant trade are the major pathways for squamate introductions (e.g., Perella & Behm 2020). Estimates of the goods produced, and volume traded from within native ranges might better explain introduction and possibly extinction-prone status (Tingley et al. 2016).

Consistent with observations in other taxa (e.g., Pacifici et al. 2020), we found that the native range geography of squamates influences both introduction and extinction status. Introduced squamates generally had larger and edgier native ranges, indicative of spatially spread out and disjoint native distributions (Fig. 3B). In contrast, extinction-prone species had smaller and more compact native ranges. The IUCN often uses convex hulls to delineate ranges. This method can cause smaller ranges to be more circular and less edgy, and species with smaller ranges are more likely to be extinction-prone (IUCN 2021). In contrast, any species that has a large and edgier native range is more likely to encounter humans and be introduced by hitchhiking along transportation networks. Edgier ranges indicate abutment to coastlines and river valleys where most of humanity resides (Small & Nicholls 2003). Further, ancient colonization explains recent naturalizations of anole lizards and edgier ranges are also indicative of disjoint distributions derived from ancient long-distance natural colonization events (Poe et al. 2011).

The complexity of past and present climatic variation over which the macroevolutionary process has played out to determine the geography and ecology of Western Hemisphere squamate species says nothing about their status today. Climatic niche differentiation—an area standardized metric of the variation in climatic conditions within ranges—did not explain status. Evolutionary range expansion rate—an age standardized metric of how much area species of a genus have expanded into—also did not explain status. Similarly, current climatic conditions of native ranges had no effect on status. Neither did genus age. These negligible effects were unexpected. Measures of climate and evolutionary age do explain native range biodiversity for vertebrates globally, including Western Hemisphere squamates (e.g., Title and Burns 2015; Marin et al. 2018; Wiens et al. 2019). Thus, anthropogenetic changes to ranges are not influenced by current climate conditions, and how species ancestors adapted and responded to past climates. In the Anthropocene, there seems a decoupling of species macroevolutionary range histories from how species ranges are changing today.

Functional traits explained status. Squamate species with larger clutches—meaning those with higher reproductive potential—were often introduced. This result is consistent with theory on introduction establishment and vulnerability to stochastic events (Mahoney et al. 2015, Allen et al. 2017). Unexpectedly, the reverse was not found. Extinction-prone species did not have smaller clutch size. Viviparous species were neither less-prone to extinction nor more likely to be introduced. The effect of clutch size on introductions is likely accentuated by the pet and wildlife trade (Li et al. 2023). Pet breeders focus on species who reproduce more, and pet introductions are a major invasion pathway for squamates (Perella & Behm 2020; Stringham & Lockwood 2018). Introduced squamates were more herbivorous and omnivorous than extinction prone squamates, which were small and carnivores (Fig. 3B). The positive association between non-carnivores and introduced status is congruent with a global study on reptiles that found herbivores most likely to establish introduced populations (Mahoney et al. 2015), theoretical work that indicates diet generalism predicts invasion success (e.g., Romanuk et al. 2009), and studies of other taxa that find that species with range expansions often have generalist diets (e.g., Pacifici et al. 2020). For extinction-prone species, however, previous work has found that it is the large, herbivorous vertebrates most at risk of extinction, and for reptiles specifically it is the large, herbivorous turtles most at risk (Mahoney et al. 2015, Tingley et al. 2016, Atwood et al. 2020, Senior et al. 2021). Thus, there is variation in the extinction-size relationship across vertebrate clades. For squamates, those of small body size may be particularly sensitive to temperature shifts (Herczeg et al. 2007) making them vulnerable to human-caused microhabitat changes caused by land development (Jesse et al. 2018). Due to squamate invasion pathways, it is unsurprising that body size did not affect introduced status. Large squamates are often introduced as pets and smaller squamates introduced via the live plant trade (Powell et al. 2011). However, there is also likely variation in the relationship between squamate body size and introduction across clades (e.g., Latella et al. 2011). More work is needed to tease apart how specific ecological traits of different clades interact with specific invasion pathways and extinction drivers.

### Opposite ends of the same trait continua

The trait continua that explained both introduced and extinction-prone status all exhibited an opposite response (Fig 3B). While human footprint in native ranges increased both introduced and extinction-prone status, no geographic, evolutionary, or ecological trait explained statuses similarly. Two traits (clutch size, body size) explained only one of the statuses. Eight out of the 13 traits we tested (ca. 60%) exhibited opposite responses that were either significant (herbivorus diet, omnivorous diet, range shape, range area) or weak (body size, number of ports, genus age, viviparity). Other studies also report opposite effect sizes for 50-73% of species-level predictors in animal taxa (Jetschke & Strayer 2008, Blackburn & Jetschke 2009, Tingley et al. 2016). The support for the opposite ends hypothesis seems to find even stronger support in plant studies (Bradshaw et al. 2008, Pandit et al. 2011), although due to their different methodology, the strength of support for the hypothesis is difficult to compare among studies. We suggest a standardized metanalysis on existing studies to test if introduced and extinction-prone species do generally fall at opposite ends of the same trait continua. For Western Hemisphere squamates, there were strong positive associations of status with insularity and human footprint (Fig 3C). Therefore, dissimilarities in species characteristics alone do not provide a full understanding of what makes a species introduced or extinct. A tiered model is needed, in which the level of human impact in the native range of a species determines the likelihood of distributional change, while its ecological and geographic traits determine if the species will expand or contract.

### Implications for insular biodiversity

Islands have always served as important steppingstones for overwater dispersal and locations from which many colonizer species originated (e.g., Harbaugh et al. 2009). However, colonization and extinction rates greatly exceed historic background rates of species loss and gain. For instance, in Hawaii, the colonization rate before human settlement has been estimated at 0.03 species per 1000 years, increasing to 20 species per 1000 years with the arrival of Polynesians and 20000 species per 1000 years over the last two centuries (Ricciardi 2007). While oceanic islands are generally more susceptible to exotic invasion than mainland regions (Dawson et al. 2017), the level of colonization pressure on Hawaii in the last centuries is likely an extreme example.

The current rate of species introductions currently surpasses the rate of species extinctions (Ellis et al. 2012). In our dataset, only eight of 483 extinction-prone species were already considered extinct, while a total of 21% of species (483 out of 2268) were threatened with extinction based on IUCN status. This threat level approximates results for all vertebrates, of which 18% are at risk of extinction (Atwood et al. 2020) and 32% of terrestrial vertebrates experience range reduction or population decline (also including low concern species) (Ceballos et al. 2017). The discrepancy between extinct and threatened species numbers indicates the existence of an extinction debt: the surplus of species that will go extinct in the foreseeable future, but still contributes to the species richness we see today.

The swift influx of colonizers and prospective loss of endemic species leads to significant taxonomic and functional homogenization especially within island biota (Longman et al. 2018). This threatens biodiversity and ecosystem functioning on a small scale, but also poses a threat to global biodiversity because of the profound risk to lose a large portion of diversity on earth. To mitigate island homogenization, screening, biosecurity measures, and targeted protection policies must be better implemented on islands.

## Acknowledgements

None

## Appendix

**Figure S1.**
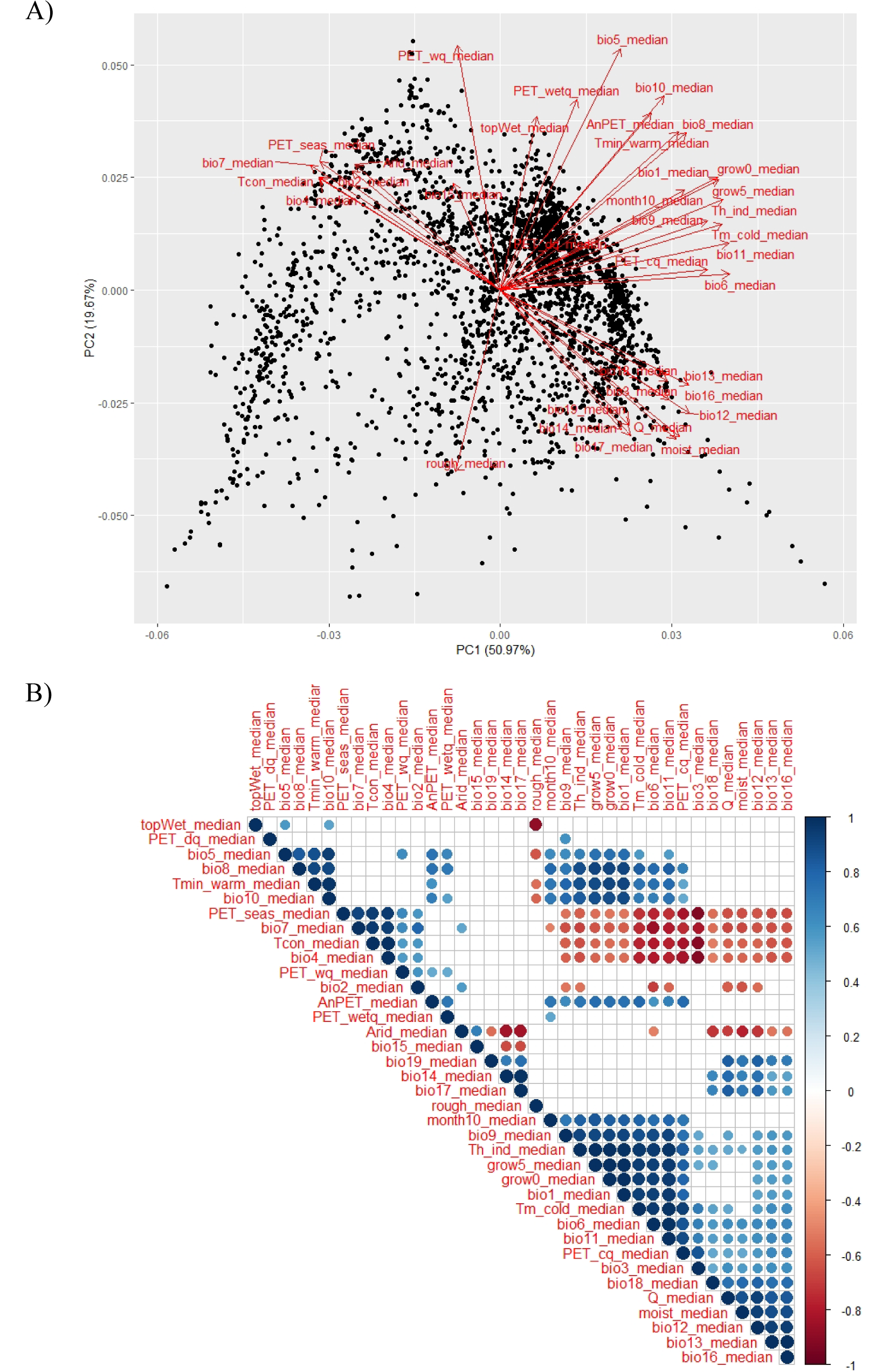
Biplot and correlation matrix including all 37 climatic variables from Woldclim (Fick et al., 2017) and ENVIREM (Title & Bemmels, 2018) databases, which were included a scaled Principal Component Analysis (PCA). From this PCA, two environmental PC axes were derived and included in further analysis. A) Biplot of axis 1 (PC1) and axis 2 (PC2) of a PCAthat shows the variation among the median values of 37 climatic variables in 3061 species ranges. 1 point is 1 species range. PC1 aligns with variables indicating the level of seasonality within a species range (e.g., temperature seasonality; continentality and temperature range, negatively correlated to temperature in the coldest month). PC2 aligns with variables associated with elevational differences (e.g., terrain roughness index; SAGA-GIS topographic wetness index). We inverted both axes compared to the rotation depicted here. B) Correlation matrix of 37 WorldClim and ENVIREM variables for which the median per species range has been calculated. The variables are ordered to maximize congruence. Correlation coefficients > −0.5 and < 0.5 were excluded from this figure for clarity. Dot size indicates the strength of the correlations; the larger the dot, the closer the correlation coefficient is to 1 or –1. Blue dots indicate a positive correlation and red dots indicate a negative correlation among variables, and thus a parallel or opposite direction of the arrows in figure S1A, respectively.

**Figure S2.**
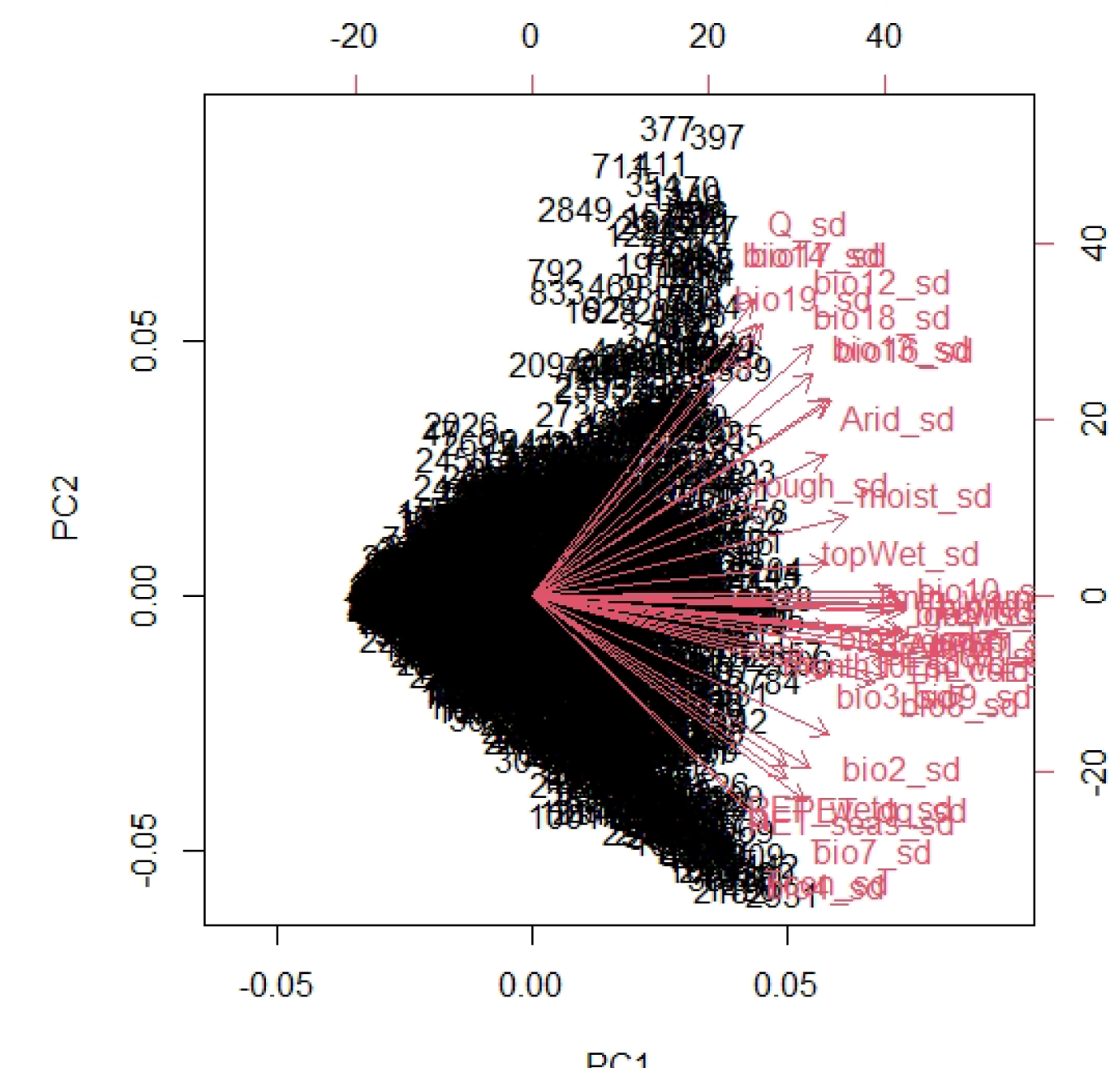
Biplot of WorldClim and ENVIREM climatic variables included in the climatic niche differentiation variable. The standard deviations of 37 climatic variables within 3056 species ranges were included in a principal component analysis (PCA). 1 point is 1 species range. PC1 aligns with all climatic variables, explaining a total of 63% of the overall variation. This means that if a standard deviation of for instance a temperature-based variable is high in a species range (i.e., a species experiences a variety of temperatures), also the standard deviations of other climatic variables increase in the same direction. And when the climatic standard deviations within a species range are high, the PC1 value for that species is also high. As it is expected that larger species ranges have more variable climates, PC1 was regressed against range size, taking the residuals of that relationship (r = 0.65, t_3054_ = 47.45, *P* < 0.001) as value for disproportional climatic niche differentiation/uniformity.

**Figure S3.**
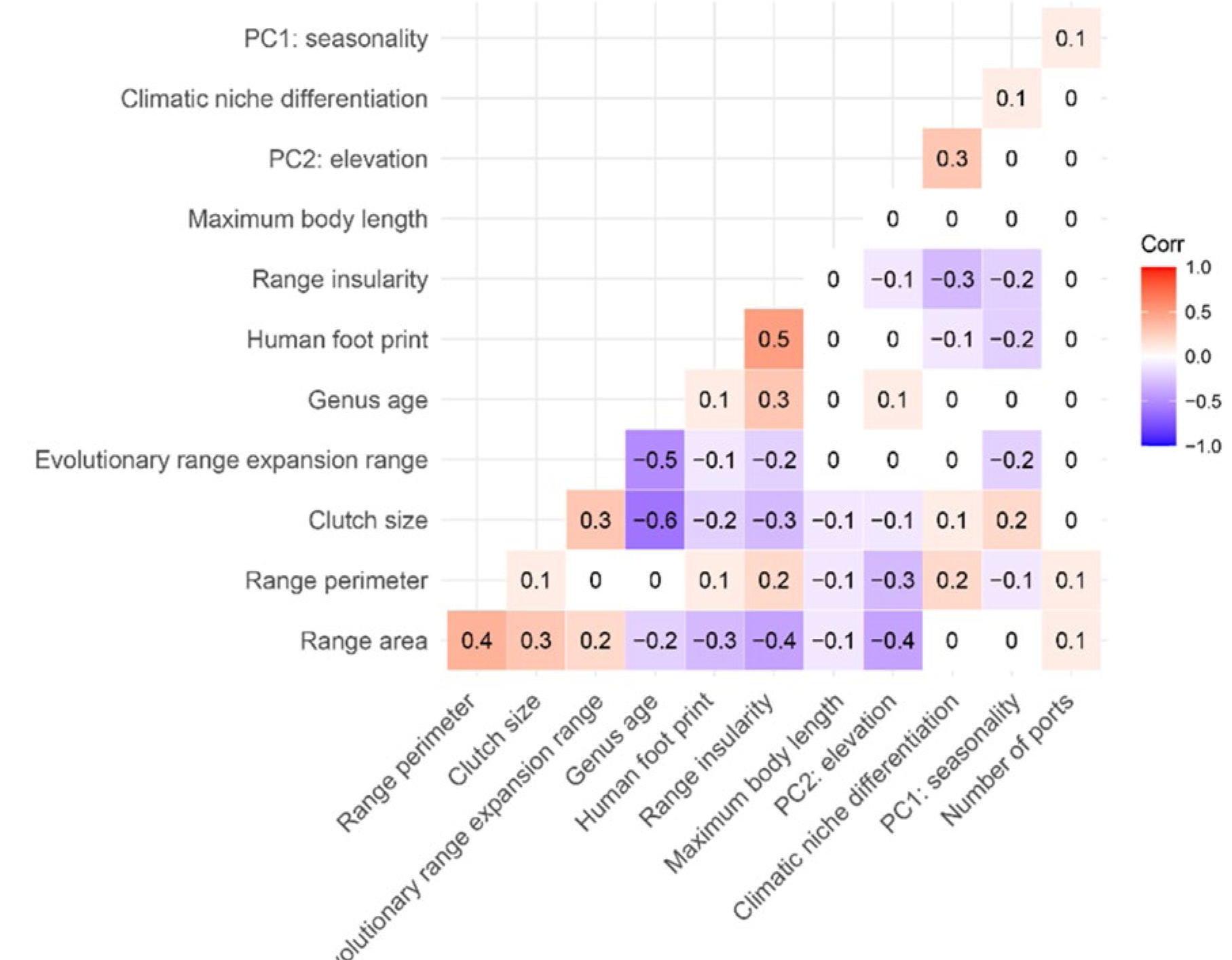
Correlations among predictor variables that were included in phylogenetic generalized linear models. Correlation matrix including all predictor variables. The brighter the color, the stronger the correlation between variables (i.e., closer to 1 or –1). Blue blocks indicate a negative relationship and red blocks indicate a positive relationship. The relatively low correlation coefficients among predictor variables indicate that there are no issues with collinear relationships that bias our model outputs (also indicated by a relatively low Variance Inflation Factor of <3, see main text)

**Table S1.**
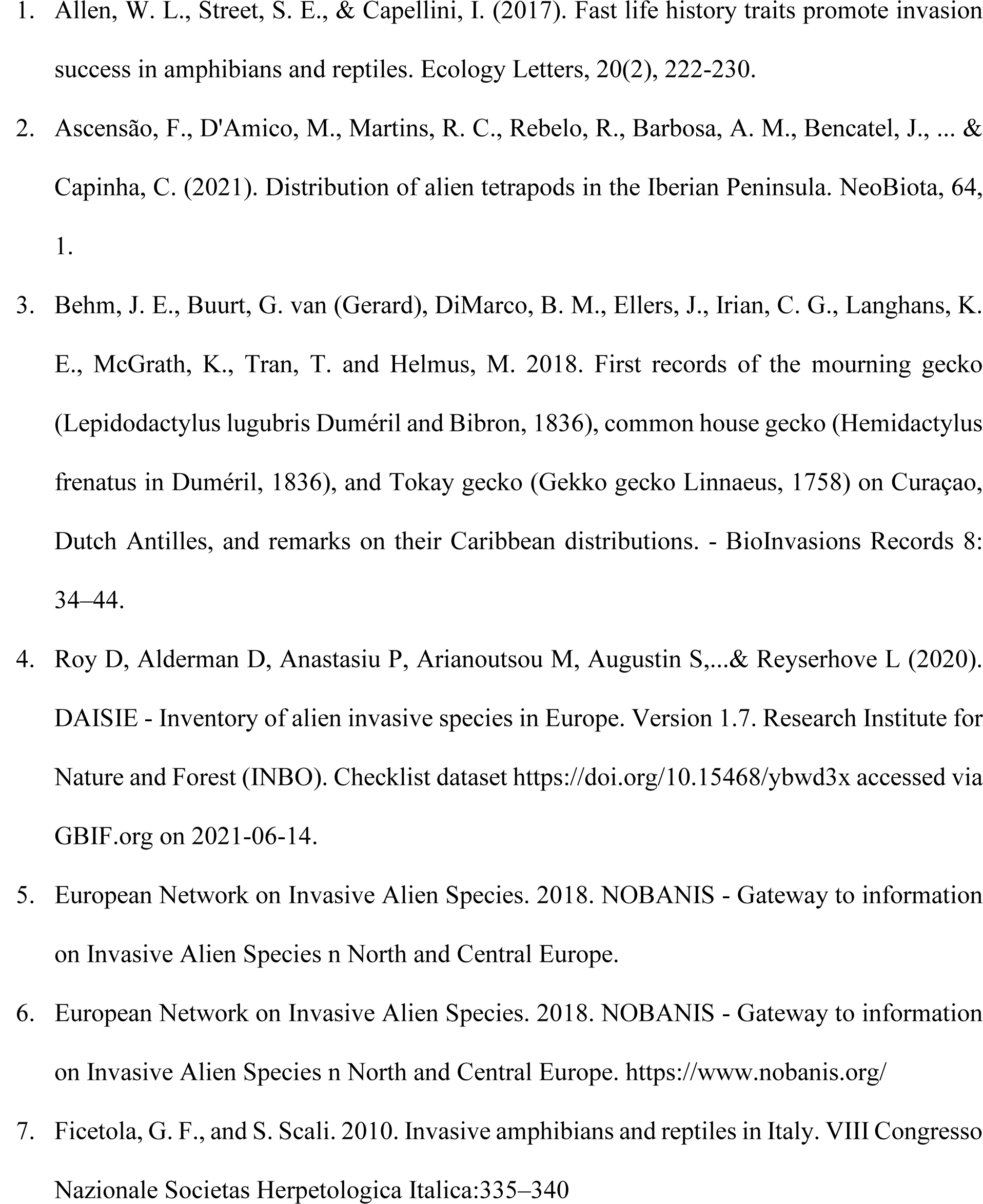
list of references used to estimate the introduction status of Western Hemisphere squamates.

**Table S2.**
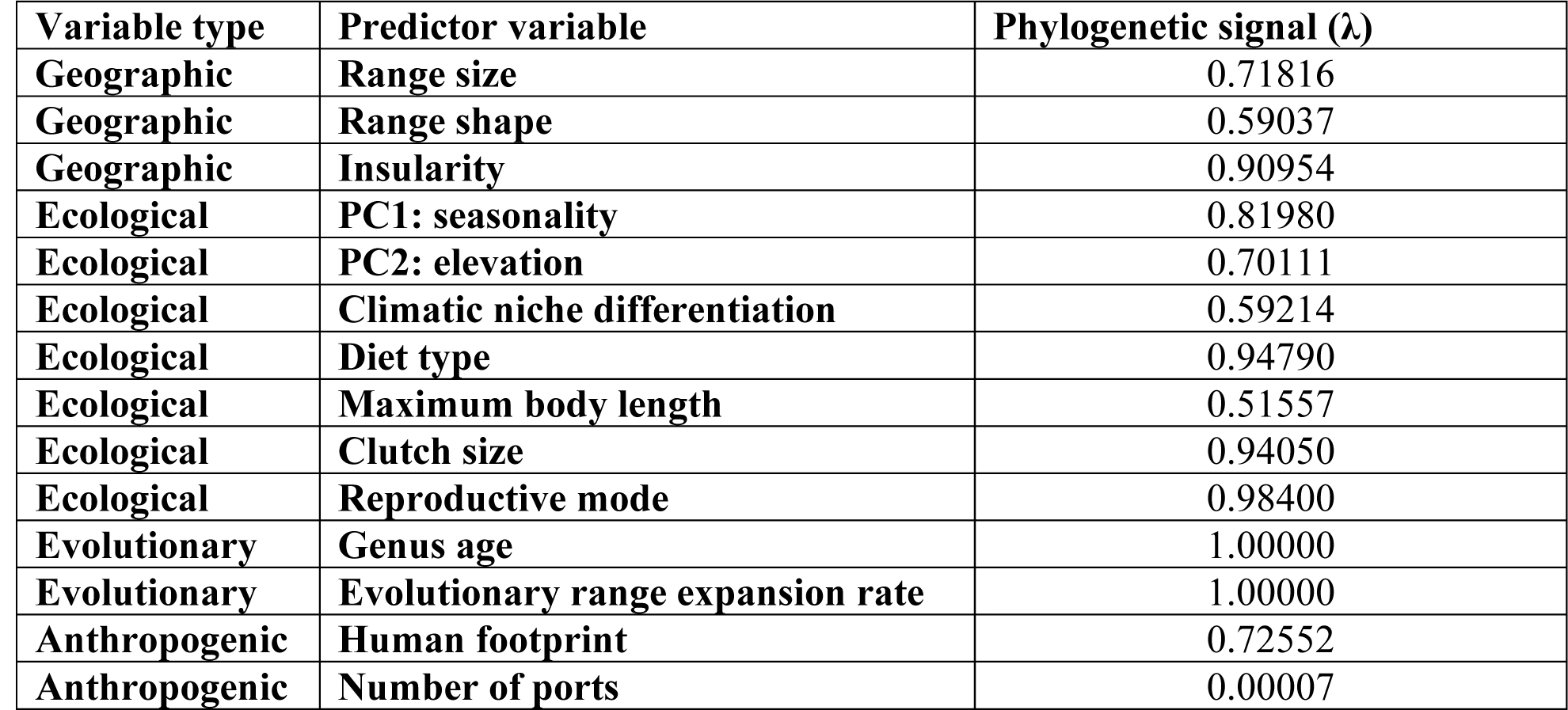
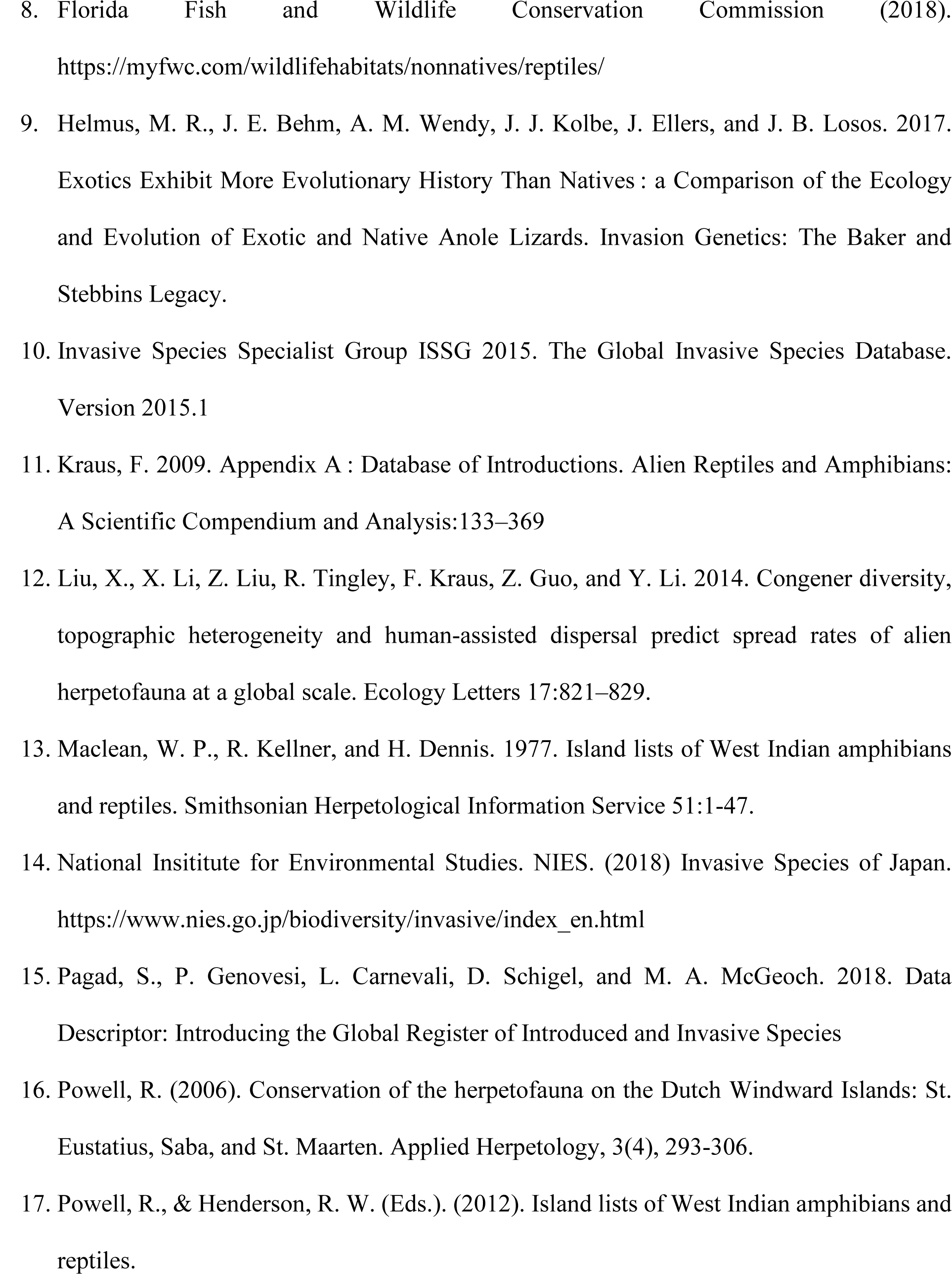
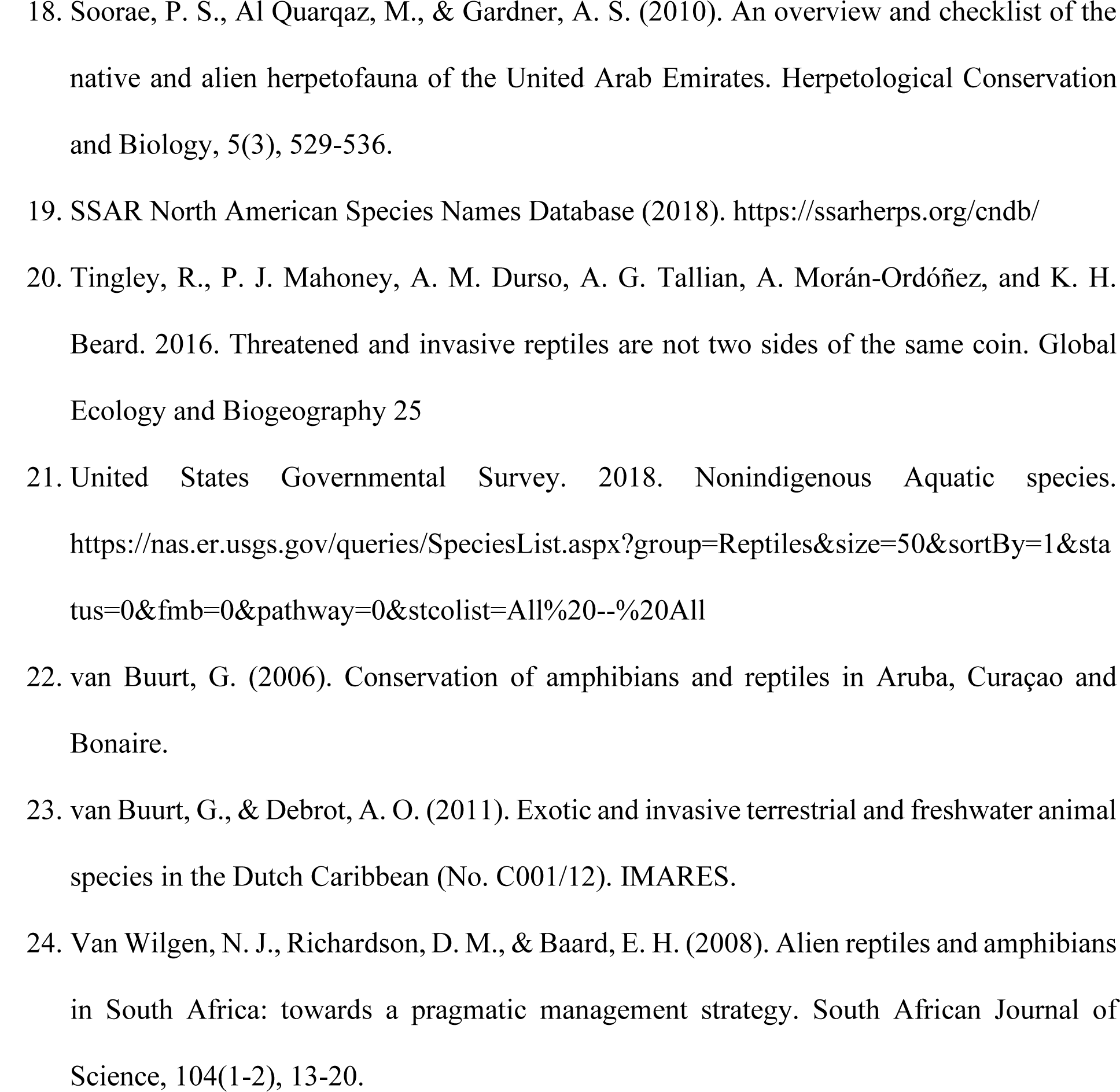
Phylogenetic signal of predictor variables. Phylogenetic signal of discrete and continuous predictor variables. Phylogenetic signal is defined as Lamda (λ), ranging between 0 and 1, the higher the value, the stronger the signal (“fitContinuous” and “fitDiscrete” functions in *geiger* R package). We expected phylogenetic signal for some variables: Clutch size, maximum body length, diet type, and reproductive mode were phylogenetically imputed (see Rapacciuolo et al. 2019 for details). Furthermore, two variables were genus-specific variables (stem age, evolutionary range expansion range), hence inherently phylogenetically clustered.

## References

Adeoba, M., S. G. Tesfamichael, and K. Yessoufou. 2019. Preserving the tree of life of the fish family Cyprinidae in Africa in the face of the ongoing extinction crisis. Genome 62:170–182.

Alcaraz, C., A. Vila-Gispert, and E. García-Berthou. 2005. Profiling invasive fish species: The importance of phylogeny and human use. Diversity and Distributions 11:289–298.

Allen, W. L., S. E. Street, and I. Capellini. 2017. Fast life history traits promote invasion success in amphibians and reptiles. Ecology Letters 2:222–230

Alroy, J. 2015. Current extinction rates of reptiles and amphibians. Proceedings of the National Academy of Sciences of the United States of America 112:13003–13008.

Amiel, J. J., R. Tingley, and R. Shine. 2011. Smart moves: Effects of relative Brain size on Establishment success of invasive amphibians and reptiles. PLoS ONE 6:e18277.

Arbetman, M. P., G. Gleiser, C. L. Morales, P. Williams, and M. A. Aizen. 2017. Global decline of bumblebees is phylogenetically structured and inversely related to species range size and pathogen incidence. Proceedings of the Royal Society B: Biological Sciences 284:20170204.

Atwood, T. B., S. A. Valentine, E. Hammill, D. J. McCauley, E. M. P. Madin, K. H. Beard, and W. D. Pearse. 2020. Herbivores at the highest risk of extinction among mammals, birds, and reptiles. Science Advances 6:eabb8458.

Behm, J., Busala, G. and Helmus, M. 2022. First records of three new lizard species and a range expansion of a fourth lizard species introduced to Aruba. BioInvasions Records 11:296–306.

Blackburn, T. M., and R. P. Duncan. 2001. Establishment patterns of exotic birds are constrained by non-random patterns in introduction. Journal of Biogeography 28:927–939.

Blackburn, T. M., and J. M. Jeschke. 2009. Invasion success and threat status: Two sides of a different coin? Ecography 32:83–88.

Böhm, M., B. Collen, J. E. M. Baillie, P. Bowles, J. Chanson, N. Cox, G. Hammerson, M. Hoffmann, S. R. Livingstone, M. Ram, A. G. J. Rhodin, S. N. Stuart, P. P. van Dijk, B. E. Young, L. E. Afuang, A. Aghasyan, A. García, C. Aguilar, R. Ajtic, F. Akarsu, L. R. V. Alencar, A. Allison, N. Ananjeva, S. Anderson, C. Andrén, D. Ariano-Sánchez, J. C. Arredondo, M. Auliya, C. C. Austin, A. Avci, P. J. Baker, A. F. Barreto-Lima, C. L. Barrio-Amorós, D. Basu, M. F. Bates, A. Batistella, A. Bauer, D. Bennett, W. Böhme, D. Broadley, R. Brown, J. Burgess, A. Captain, S. Carreira, M. del R. Castañeda, F. Castro, A. Catenazzi, J. R. Cedeño-Vázquez, D. G. Chapple, M. Cheylan, D. F. Cisneros-Heredia, D. Cogalniceanu, H. Cogger, C. Corti, G. C. Costa, P. J. Couper, T. Courtney, J. Crnobrnja-Isailovic, P. A. Crochet, B. Crother, F. Cruz, J. C. Daltry, R. J. R. Daniels, I. Das, A. de Silva, A. C. Diesmos, L. Dirksen, T. M. Doan, C. K. Dodd, J. S. Doody, M. E. Dorcas, J. Duarte de Barros Filho, V. T. Egan, E. H. El Mouden, D. Embert, R. E. Espinoza, A. Fallabrino, X. Feng, Z. J. Feng, L. Fitzgerald, O. Flores-Villela, F. G. R. França, D. Frost, H. Gadsden, T. Gamble, S. R. Ganesh, M. A. Garcia, J. E. García-Pérez, J. Gatus, M. Gaulke, P. Geniez, A. Georges, J. Gerlach, S. Goldberg, J. C. T. Gonzalez, D. J. Gower, T. Grant, E. Greenbaum, C. Grieco, P. Guo, A. M. Hamilton, K. Hare, S. B. Hedges, N. Heideman, C. Hilton-Taylor, R. Hitchmough, B. Hollingsworth, M. Hutchinson, I. Ineich, J. Iverson, F. M. Jaksic, R. Jenkins, U. Joger, R. Jose, Y. Kaska, U. Kaya, J. S. Keogh, G. Köhler, G. Kuchling, Y. Kumlutaş, A. Kwet, E. La Marca, W. Lamar, A. Lane, B. Lardner, C. Latta, G. Latta, M. Lau, P. Lavin, D. Lawson, M. LeBreton, E. Lehr, D. Limpus, N. Lipczynski, A. S. Lobo, M. A. López-Luna, L. Luiselli, V. Lukoschek, M. Lundberg, P. Lymberakis, R. Macey, W. E. Magnusson, D. L. Mahler, A. Malhotra, J. Mariaux, B. Maritz, O. A. V. Marques, R. Márquez, M. Martins, G. Masterson, J. A. Mateo, R. Mathew, N. Mathews, G. Mayer, J. R. McCranie, G. J. Measey, F. Mendoza-Quijano, M. Menegon, S. Métrailler, D. A. Milton, C. Montgomery, S. A. A. Morato, T. Mott, A. Muñoz-Alonso, J. Murphy, T. Q. Nguyen, G. Nilson, C. Nogueira, H. Núñez, N. Orlov, H. Ota, J. Ottenwalder, T. Papenfuss, S. Pasachnik, P. Passos, O. S. G. Pauwels, N. Pérez-Buitrago, V. Pérez-Mellado, E. R. Pianka, J. Pleguezuelos, C. Pollock, P. Ponce-Campos, R. Powell, F. Pupin, G. E. Quintero Díaz, R. Radder, J. Ramer, A. R. Rasmussen, C. Raxworthy, R. Reynolds, N. Richman, E. L. Rico, E. Riservato, G. Rivas, P. L. B. da Rocha, M. O. Rödel, L. Rodríguez Schettino, W. M. Roosenburg, J. P. Ross, R. Sadek, K. Sanders, G. Santos-Barrera, H. H. Schleich, B. R. Schmidt, A. Schmitz, M. Sharifi, G. Shea, H. T. Shi, R. Shine, R. Sindaco, T. Slimani, R. Somaweera, S. Spawls, P. Stafford, R. Stuebing, S. Sweet, E. Sy, H. J. Temple, M. F. Tognelli, K. Tolley, P. J. Tolson, B. Tuniyev, S. Tuniyev, N. üzüm, G. van Buurt, M. Van Sluys, A. Velasco, M. Vences, M. Veselý, S. Vinke, T. Vinke, G. Vogel, M. Vogrin, R. C. Vogt, O. R. Wearn, Y. L. Werner, M. J. Whiting, T. Wiewandt, J. Wilkinson, B. Wilson, S. Wren, T. Zamin, K. Zhou, and G. Zug. 2013. The conservation status of the world’s reptiles. Biological Conservation 157:372–385.

Böhm, M., R. Williams, H. R. Bramhall, K. M. Mcmillan, A. D. Davidson, A. Garcia, L. M. Bland, J. Bielby, and B. Collen. 2016. Correlates of extinction risk in squamate reptiles: The relative importance of biology, geography, threat and range size. Global Ecology and Biogeography 25:391–405.

Bradshaw, C. J. A., X. Giam, H. T. W. Tan, B. W. Brook, and N. S. Sodhi. 2008. Threat or invasive status in legumes is related to opposite extremes of the same ecological and life-history attributes. Journal of Ecology 96:869–883.

Cadotte, M. W., M. A. Hamilton, and B. R. Murray. 2009. Phylogenetic relatedness and plant invader success across two spatial scales. Diversity and Distributions 15:481–488.

Capinha, C., F. Essl, H. Seebens, D. Moser, and H. M. Pereira. 2015. The dispersal of alien species redefines biogeography in the Anthropocene. Science 348:1248–1251.

Ceballos, G., P. R. Ehrlich, and R. Dirzo. 2017. Biological annihilation via the ongoing sixth mass extinction signaled by vertebrate population losses and declines. Proceedings of the National Academy of Sciences of the United States of America 114:E6089–E6096.

Chichorro, F., Juslén, A. & Cardoso, P. (2019). A review of the relation between species traits and extinction risk. Biological Conservation, 237:220–229.

Colautti, R. I., and H. I. MacIsaac. 2004. A neutral terminology to define “invasive” species. Diversity and Distributions 10:135–141.

Cowie, R. H., and B. S. Holland. 2006. Dispersal is fundamental to biogeography and the evolution of biodiversity on oceanic islands. 33:193–198.

Cox, N., Young, B.E., Bowles, P., Fernandez, M., Marin, J., Rapacciuolo, G., et al. (2022). A global reptile assessment highlights shared conservation needs of tetrapods. Nature, 605: 285–290.

Davies, T. J., G. F. Smith, D. U. Bellstedt, J. S. Boatwright, B. Bytebier, R. M. Cowling, F. Forest, L. J. Harmon, A. M. Muasya, B. D. Schrire, Y. Steenkamp, M. van der Bank, and V. Savolainen. 2011. Extinction risk and diversification are linked in a plant biodiversity hotspot. PLoS Biology 9: e1000620.

Dawson, W., Moser, D., Van Kleunen, M., Kreft, H., Pergl, J., Pyšek, P., Weigelt, P., Winter, M., Lenzner, B., Blackburn, T.M. and Dyer, E.E., 2017. Global hotspots and correlates of alien species richness across taxonomic groups. Nature Ecology & Evolution 1:1–7.

Ellis, E. C., E. C. Antill, and H. Kreft. 2012. All is not loss: plant biodiversity in the Anthropocene. PloS ONE 7:e30535.

Fernández-Palacios, J.M., Kreft, H., Irl, S.D.H., Norder, S., Ah-Peng, C., Borges, P.A.V., et al. 2021. Scientists’ warning – The outstanding biodiversity of islands is in peril. Global Ecology and Conservation 31:e01847.

Fick, S. E., and R. J. Hijmans. 2017. WorldClim 2: new 1-km spatial resolution climate surfaces for global land areas. International Journal of Climatology 37:4302–4315.

Fordham, D. A., and B. W. Brook. 2010. Why tropical island endemics are acutely susceptible to global change. Biodiversity and Conservation 19:392–342.

Frazier, A.E. & Kedron, P. (2017). Landscape metrics: past progress and future directions. Current Landscape Ecology Reports. 2:63–72.

Fritz, S. A., and A. Purvis. 2010. Selectivity in mammalian extinction risk and threat types: A new measure of phylogenetic signal strength in binary traits. Conservation Biology 24:1042–1051.

GADM. 2018. Database of Global Administrative Areas, version 3.6. The University of California, Berkley.

Gleditsch, J. M., Behm, J. E., Ellers, J., Jesse, W. A., & Helmus, M. R. 2023. Contemporizing island biogeography theory with anthropogenic drivers of species richness. Global Ecology and Biogeography 32:233–249.

Harbaugh, D. T., W. L. Wagner, G. J. Allan, and E. A. Zimmer. 2009. The Hawaiian Archipelago is a stepping stone for dispersal in the Pacific: an example from the plant genus Melicope (Rutaceae). Journal of Biogeography 36:230–241.

Hedges, S. B. 2006. An overview of the evolution and conservation of West Indian amphibians and reptiles. Applied Herpetology 3:281–292.

Hedges, S.B. 2021. Caribherp: amphibians and reptiles of Caribbean Islands. Available online at http://www.caribherp.org/ (accessed 1 July 2021). Philadelphia, Pennsylvania: Temple University.

Helmus, M. R., J. E. Behm, A. M. Wendy, J. J. Kolbe, J. Ellers, and J. B. Losos. 2017. Exotics Exhibit More Evolutionary History Than Natives: A Comparison of the Ecology and Evolution of Exotic and Native Anole Lizards. Invasion Genetics: The Baker and Stebbins Legacy 122–138.

Herczeg, G., J. Török, and Z. Korsós. 2007. Size-dependent heating rates determine the spatial and temporal distribution of small-bodied lizards. Amphibia Reptilia 28:347–356.

Hijmans, R. J. 2019. geosphere: Spherical Trigonometry. R package version 1.5–10.

Ho, L.S.T. & Ane, C. 2014. A linear-time algorithm for Gaussian and non-Gaussian trait evolution models. Systematic Biology 63:397–408.

IPBES. 2019. Global assessment report on biodiversity and ecosystem services of the Intergovernmental Science-Policy Platform on Biodiversity and Ecosystem Services. E. S. Brondizio, J. Settele, S. Díaz, and H. T. Ngo (editors). IPBES secretariat, Bonn, Germany. 1148 pages. 10.5281/zenodo.3831673.

IPBES. 2023. Summary for Policymakers of the Thematic Assessment Report on Invasive Alien Species and their Control of the Intergovernmental Science-Policy Platform on Biodiversity and Ecosystem Services. Roy, H. E., Pauchard, A., Stoett, P., Renard Truong, T., Bacher, S., Galil, B. S., Hulme, P. E., Ikeda, T., Sankaran, K. V., McGeoch, M. A., Meyerson, L. A., Nuñez, M. A., Ordonez, A., Rahlao, S. J., Schwindt, E., Seebens, H., Sheppard, A. W., and Vandvik, V. (eds.). IPBES secretariat, Bonn, Germany. 10.5281/zenodo.7430692.

IUCN Redlist. 2021. IUCN Redlist of Threatened Species. Available from http://www.iucnredlist.org/ (accessed 1 July 2021). Gland, Switzerland: International Union for the Conservation of Nature.

Jantz, S. M., B. Barker, T. M. Brooks, L. P. Chini, Q. Huang, R. M. Moore, J. Noel, and G. C. Hurtt. 2015. Future habitat loss and extinctions driven by land-use change in biodiversity hotspots under four scenarios of climate-change mitigation. Conservation Biology 29:1122–1131.

Jeschke, J. M., and D. L. Strayer. 2008. Are threat status and invasion success two sides of the same coin? Ecography 31:124–130.

Jesse, W. A. M., J. E. Behm, M. R. Helmus, and J. Ellers. 2018. Human land use promotes the abundance and diversity of exotic species on Caribbean islands. Global Change Biology 24:4784–4796.

Kier, G., H. Kreft, T. M. Lee, W. Jetz, P. L. Ibisch, C. Nowicki, J. Mutke. and W. Barthlott. 2009. A global assessment of endemism and species richness across island and mainland regions. Proceedings of the National Academy of Sciences, 106: 9322–9327.

Kotiaho, J. S., V. Kaitala, A. Komonen, and J. Päivinen. 2005. Predicting the risk of extinction from shared ecological characteristics. Proceedings of the National Academy of Sciences of the United States of America 102:1963–1967.

Krummel, J.R., Gardner, R.H., Sugihara, G., O’Neill, R.V. & Coleman, P.R. (1987). Landscape patterns in a disturbed environment. Oikos 48:321–324.

Kupfer, J.A. (2012). Landscape ecology and biogeography: Rethinking landscape metrics in a post-FRAGSTATS landscape. Progress in Physical Geography: Earth and Environment, 36:400–420.

Kumar, S., G. Stecher, M. Suleski, and S. B. Hedges. 2017. TimeTree: A Resource for Timelines, Timetrees, and Divergence Times. Molecular biology and evolution 34:1812–1819.

Larson, E. R., and J. D. Olden. 2010. Latent extinction and invasion risk of crayfishes in the southeastern United States. Conservation Biology 24:1099–1110.

Latella, I. M., S. Poe, and J. T. Giermakowski. 2011. Traits associated with naturalization in Anolis lizards: Comparison of morphological, distributional, anthropogenic, and phylogenetic models. Biological Invasions 13:845–856.

Li, Y., Blackburn, T. M., Luo, Z., Song, T., Watters, F., Li, W., Deng, T., Luo, Z., Li, Y., Du, J., Niu, M., Zhang, J., Zhang, J., Yang, J. and Wang, S. 2023. Quantifying global colonization pressures of alien vertebrates from wildlife trade. Nat Commun 14: 7914.

Liu, C., He, D., Chen, Y. and Olden, J. D. 2017. Species invasions threaten the antiquity of China’s freshwater fish fauna. Diversity and Distributions 23: 556–566.

Liu, X., X. Li, Z. Liu, R. Tingley, F. Kraus, Z. Guo, and Y. Li. 2014. Congener diversity, topographic heterogeneity and human-assisted dispersal predict spread rates of alien herpetofauna at a global scale. Ecology Letters 17:821–829.

Longman, E. K., K. Rosenblad, and D. F. Sax. 2018. Extreme homogenization: The past, present and future of mammal assemblages on islands. Global Ecology and Biogeography 27:77–95.

Losos, J. B. 2011. Lizards in an Evolutionary Tree: Ecology and Adaptive Radiation of Anoles. University of California Press.

Loza, M. I., I. Jiménez, P. M. Jørgensen, G. Arellano, M. J. Macía, V. W. Torrez, and R. E. Ricklefs. 2017. Phylogenetic patterns of rarity in a regional species pool of tropical woody plants. Global Ecology and Biogeography 26:1043–1054.

Mahler, D. L., T. Ingram, L. J. Revell, and J. B. Losos. 2013. Exceptional convergence on the macroevolutionary landscape in island lizard radiations. Science 341:292–295.

Mahler, D. L., L. J. Revell, R. E. Glor, and J. B. Losos. 2010. Ecological opportunity and the rate of morphological evolution in the diversification of greater Antillean anoles. Evolution 64:2731–2745.

Mahoney, P. J., K. H. Beard, A. M. Durso, A. G. Tallian, A. L. Long, R. J. Kindermann, N. E. Nolan, D. Kinka, and H. E. Mohn. 2015. Introduction effort, climate matching and species traits as predictors of global establishment success in non-native reptiles. Diversity and Distributions 21:64–74.

MacArthur, R. H. and E.O. Wilson. 1967. Island biogeography. Princeton, USA.

Marin, J., G. Rapacciuolo, G. C. Costa, C. H. Graham, T. M. Brooks, B. E. Young, V. C. Radeloff, J. E. Behm, M. R. Helmus, and S. B. Hedges. 2018. Evolutionary time drives global tetrapod diversity. Proceedings of the Royal Society B: Biological Sciences 285:20172378.

Marino, C. & Bellard, C. 2023. When origin, reproduction ability and diet define the role of birds in invasions. Proc. R. Soc. B. 290:20230196.

Mittermeier, R. A., Turner, W. R., Larsen, F. W., Brooks, T. M., & Gascon, C. (2011). Global biodiversity conservation: the critical role of hotspots. In Biodiversity hotspots: distribution and protection of conservation priority areas (pp. 3-22). Berlin, Heidelberg: Springer Berlin Heidelberg.

Monaco, C. J., C. J. A. Bradshaw, D. J. Booth, B. M. Gillanders, D. S. Schoeman, and I. Nagelkerken. 2020. Dietary generalism accelerates arrival and persistence of coral-reef fishes in their novel ranges under climate change. Global Change Biology 26:5564–5573.

Myers, N., R. A. Mittermeler, C. G. Mittermeler, G. A. B. Da Fonseca, and J. Kent. 2000. Biodiversity hotspots for conservation priorities. Nature 403:853–858.

National Geospatial-Intelligence Agency. 2017. World Port Index. https://msi.nga.mil/Publications/WPI.

Orme, D., Freckleton, R., Thomas, G., Petzoldt, T., Fritz, S., Isaac, N., & Pearse, W. 2013. The caper package: comparative analysis of phylogenetics and evolution in R. R package version 5:1–36.

Pandit, M. K., M. J. O. Pocock, and W. E. Kunin. 2011. Ploidy influences rarity and invasiveness in plants. Journal of Ecology 99:1108–1115.

Park, D. S., and D. Potter. 2015. Why close relatives make bad neighbours: Phylogenetic conservatism in niche preferences and dispersal disproves Darwin’s naturalization hypothesis in the thistle tribe. Molecular Ecology 24:3181–3193.

Patton, D. R. 1975. A diversity index for quantifying habital “edge”. Wildlife Society Bulletin (1973–2006). 3:171-173.

Perella, C. D. and Behm, J. E. 2020. Understanding the spread and impact of exotic geckos in the greater Caribbean region. Biodivers Conserv 29:1109–1134.

Perry, G., Razi Dmi’el, & Lazell, J. 1999. Evaporative Water Loss in Insular Populations of the Anolis cristatellus Group (Reptilia: Sauria) in the British Virgin Islands II: The Effects of Drought. Biotropica 31:337–343.

Pigot, A. L., E. E. Dyer, D. W. Redding, P. Cassey, G. H. Thomas, and T. M. Blackburn. 2018. Species invasions and the phylogenetic signal in geographical range size. 27:1080–1092.

Poe, S., J. T. Giermakowski, I. Latella, E. W. Schaad, E. P. Hulebak, and M. J. Ryan. 2011. Ancient colonization predicts recent naturalization in anolis lizards. Evolution 65:1195–1202.

Powell, R., Henderson, R. W., Farmer, M. C., Breuil, M., Echternacht, A. C., Van Buurt, G.,…& Perry, G. (2011). Introduced amphibians and reptiles in the Greater Caribbean: Patterns and conservation implications. In Conservation of Caribbean Island herpetofaunas volume 1: conservation biology and the wider Caribbean (pp. 63-143). Brill.

Pyšek, P., J. Pergl, F. Essl, B. Lenzner, W. Dawson, H. Kreft, P. Weigelt, M. Winter, J. Kartesz, M. Nishino, L. A. Antonova, J. F. Barcelona, F. J. Cabezas, D. Cárdenas, J. Cárdenas-Toro, N. Castaño, E. Chacón, C. Chatelain, S. Dullinger, A. L. Ebel, E. Figueiredo, N. Fuentes, P. Genovesi, Q. J. Groom, L. Henderson, Inderjit, A. Kupriyanov, S. Masciadri, N. Maurel, J. Meerman, O. Morozova, D. Moser, D. Nickrent, P. M. Nowak, S. Pagad, A. Patzelt, P. B. Pelser, H. Seebens, W. S. Shu, J. Thomas, M. Velayos, E. Weber, J. J. Wieringa, M. P. Baptiste, and M. Van Kleunen. 2017. Naturalized alien flora of the world: Species diversity, taxonomic and phylogenetic patterns, geographic distribution and global hotspots of plant invasion. Preslia 89:203-274.

Rapacciuolo, G., C. H. Graham, J. Marin, J. E. Behm, G. C. Costa, S. B. Hedges, M. R. Helmus, V. C. Radeloff, B. E. Young, and T. M. Brooks. 2019. Species diversity as a surrogate for conservation of phylogenetic and functional diversity in terrestrial vertebrates across the Americas. Nature Ecology and Evolution 3:53–61.

Ricciardi, A. 2007. Are modern biological invasions an unprecedented form of global change? Conservation Biology. 21:329:336

Romanuk, T. N., Y. Zhou, U. Brose, E. L. Berlow, R. J. Williams, and N. D. Martinez. 2009. Predicting invasion success in complex ecological networks. Philosophical Transactions of the Royal Society B: Biological Sciences 364:1743–1754.

Salazar JC, Del Rosario Castañeda M, Londoño GA, Bodensteiner BL, Muñoz MM. 2019. Physiological evolution during adaptive radiation: A test of the island effect in Anolis lizards. Evolution 73:1241-1252.

Sax, D. F., and S. D. Gaines. 2008. Species invasions and extinction: The future of native biodiversity on islands. Proceedings of the National Academy of Sciences of the United States of America 105:11490–11497.

Sayol, F., Cooke, R.S.C., Pigot, A.L., Blackburn, T.M., Tobias, J.A., Steinbauer, M.J., et al. 2021. Loss of functional diversity through anthropogenic extinctions of island birds is not offset by biotic invasions. Science Advances, 7:eabj5790.

Schmidt, J. P., P. R. Stephens, and J. M. Drake. 2012. Two sides of the same coin Rare and pest plants native to the United States and Canada. Ecological Applications 22:1512–1525.

Schmidt, J.P., Davies, T.J. & Farrell, M.J. 2021. Opposing macroevolutionary and trait-mediated patterns of threat and naturalisation in flowering plants. Ecology Letters 24:1237–1250.

Senior, A. F., Böhm, M., Johnstone, C. P., McGee, M. D., Meiri, S., Chapple, D. G., & Tingley, R. 2021. Correlates of extinction risk in Australian squamate reptiles. Journal of Biogeography 48:2144–2152.

Small, C. & Nicholls, R.J. (2003). A Global Analysis of Human Settlement in Coastal Zones. Journal of Coastal Research 19:584–599.

Smith, K. G., and R. J. Almeida. 2020. When are extinctions simply bad luck? Rarefaction as a framework for disentangling selective and stochastic extinctions. Journal of Applied Ecology 57:101–110.

Spatz, D. R., K. M. Zilliacus, N. D. Holmes, S. H. M. Butchart, P. Genovesi, G. Ceballos, B. R. Tershy, and D. A. Croll. 2017. Globally threatened vertebrates on islands with invasive species Science Advances 3: e1603080.

Stringham, O. C. and J. L. Lockwood. 2018. Pet problems: biological and economic factors that influence the release of alien reptiles and amphibians by pet owners. Journal of Applied Ecology 55:2632–2640.

Stroud, J. T. 2021. Island species experience higher niche expansion and lower niche conservatism during invasion. Proceedings of the National Academy of Sciences 118:e2018949118.

Su, S., P. Cassey, and T. M. Blackburn. 2016. The wildlife pet trade as a driver of introduction and establishment in alien birds in Taiwan. Biological Invasions 18:215–229.

Tingley, R., P. J. Mahoney, A. M. Durso, A. G. Tallian, A. Morán-Ordóñez, and K. H. Beard. 2016. Threatened and invasive reptiles are not two sides of the same coin. Global Ecology and Biogeography 25:1050–1060.

Title, P. O., and J. B. Bemmels. 2018. ENVIREM: an expanded set of bioclimatic and topographic variables increases flexibility and improves performance of ecological niche modeling. Ecography 41:291–307.

Title, P. O., and K. J. Burns. 2015. Rates of climatic niche evolution are correlated with species richness in a large and ecologically diverse radiation of songbirds. Ecology Letters 18:433–440.

Tonini, J. F. R., Beard, K. H., Ferreira, R. B., Jetz, W., & Pyron, R. A. 2016. Fully-sampled phylogenies of squamates reveal evolutionary patterns in threat status. Biological Conservation 204:23–31.

Uetz, P., Freed, P., Aguilar, R., Reyes, F., Kudera, J. & Hošek, J. (Eds.). 2021. The Reptile Database, http://www.reptile-database.org.

Venter, O., E. W. Sanderson, A. Magrach, J. R. Allan, J. Beher, K. R. Jones, H. P. Possingham, W. F. Laurance, P. Wood, B. M. Fekete, M. A. Levy, and J. E. M. Watson. 2016. Global terrestrial Human Footprint maps for 1993 and 2009. Scientific Data 3:160067.

Vitousek, P. M., H. A. Mooney, J. Lubchenco, and J. M. Melillo. 1997. Human domination of Earth’s ecosystems. Science 277:494–499.

Waters, C.N. & Turner, S.D. 2022. Defining the onset of the Anthropocene. Science, 378, 706–708.

Wiens, J. J., Y. Litvinenko, L. Harris, and T. Jezkova. 2019. Rapid niche shifts in introduced species can be a million times faster than changes among native species and ten times faster than climate change. Journal of Biogeography 46:2115–2125.

Yessoufou, K., K. Mearns, H. O. Elansary, and G. H. Stoffberg. 2016. Assessing the phylogenetic dimension of Australian Acacia species introduced outside their native ranges. Botany Letters 163:33–39.

Young, H. S., D. J. McCauley, M. Galetti, and R. Dirzo. 2016. Patterns, Causes, and Consequences of Anthropocene Defaunation. Annual Review of Ecology, Evolution, and Systematics 47:333–358.

