## appendix for "Elevated human impact on islands increases the introduction and extinction status of native insular reptiles"

**Table S1** list of references used to estimate the introduction status of Western Hemisphere squamates.


1. Allen, W. L., Street, S. E., & Capellini, I. (2017). Fast life history traits promote invasion success in amphibians and reptiles. Ecology Letters, 20(2), 222-230.
2. Ascensão, F., D'Amico, M., Martins, R. C., Rebelo, R., Barbosa, A. M., Bencatel, J., ... & Capinha, C. (2021). Distribution of alien tetrapods in the Iberian Peninsula. NeoBiota, 64, 1.
3. Behm, J. E., Buurt, G. van (Gerard), DiMarco, B. M., Ellers, J., Irian, C. G., Langhans, K. E., McGrath, K., Tran, T. and Helmus, M. 2018. First records of the mourning gecko (Lepidodactylus lugubris Duméril and Bibron, 1836), common house gecko (Hemidactylus frenatus in Duméril, 1836), and Tokay gecko (Gekko gecko Linnaeus, 1758) on Curaçao, Dutch Antilles, and remarks on their Caribbean distributions. - BioInvasions Records 8: 34–44.
4. Roy D, Alderman D, Anastasiu P, Arianoutsou M, Augustin S,...& Reyserhove L (2020). DAISIE - Inventory of alien invasive species in Europe. Version 1.7. Research Institute for Nature and Forest (INBO). Checklist dataset https://doi.org/10.15468/ybwd3x accessed via GBIF.org on 2021-06-14.
5. European Network on Invasive Alien Species. 2018. NOBANIS - Gateway to information on Invasive Alien Species n North and Central Europe.
6. European Network on Invasive Alien Species. 2018. NOBANIS - Gateway to information on Invasive Alien Species n North and Central Europe. https://www.nobanis.org/
7. Ficetola, G. F., and S. Scali. 2010. Invasive amphibians and reptiles in Italy. VIII Congresso Nazionale Societas Herpetologica Italica:335–340

8. Florida Fish and Wildlife Conservation Commission (2018). https://myfwc.com/wildlifehabitats/nonnatives/reptiles/
9. Helmus, M. R., J. E. Behm, A. M. Wendy, J. J. Kolbe, J. Ellers, and J. B. Losos. 2017. Exotics Exhibit More Evolutionary History Than Natives : a Comparison of the Ecology and Evolution of Exotic and Native Anole Lizards. Invasion Genetics: The Baker and Stebbins Legacy.
10. Invasive Species Specialist Group ISSG 2015. The Global Invasive Species Database. Version 2015.1
11. Kraus, F. 2009. Appendix A : Database of Introductions. Alien Reptiles and Amphibians: A Scientific Compendium and Analysis:133–369
12. Liu, X., X. Li, Z. Liu, R. Tingley, F. Kraus, Z. Guo, and Y. Li. 2014. Congener diversity, topographic heterogeneity and human-assisted dispersal predict spread rates of alien herpetofauna at a global scale. Ecology Letters 17:821–829.
13. Maclean, W. P., R. Kellner, and H. Dennis. 1977. Island lists of West Indian amphibians and reptiles. Smithsonian Herpetological Information Service 51:1-47.
14. National Insititute for Environmental Studies. NIES. (2018) Invasive Species of Japan. https://www.nies.go.jp/biodiversity/invasive/index_en.html
15. Pagad, S., P. Genovesi, L. Carnevali, D. Schigel, and M. A. McGeoch. 2018. Data Descriptor: Introducing the Global Register of Introduced and Invasive Species
16. Powell, R. (2006). Conservation of the herpetofauna on the Dutch Windward Islands: St. Eustatius, Saba, and St. Maarten. Applied Herpetology, 3(4), 293-306.
17. Powell, R., & Henderson, R. W. (Eds.). (2012). Island lists of West Indian amphibians and reptiles.

18. Soorae, P. S., Al Quarqaz, M., & Gardner, A. S. (2010). An overview and checklist of the native and alien herpetofauna of the United Arab Emirates. Herpetological Conservation and Biology, 5(3), 529-536.
19. SSAR North American Species Names Database (2018). https://ssarherps.org/cndb/
20. Tingley, R., P. J. Mahoney, A. M. Durso, A. G. Tallian, A. Morán-Ordóñez, and K. H. Beard. 2016. Threatened and invasive reptiles are not two sides of the same coin. Global Ecology and Biogeography 25
21. United States Governmental Survey. 2018. Nonindigenous Aquatic species. https://nas.er.usgs.gov/queries/SpeciesList.aspx?group=Reptiles&size=50&sortBy=1&status=0&fmb=0&pathway=0&stcolist=All%20--%20All
22. van Buurt, G. (2006). Conservation of amphibians and reptiles in Aruba, Curaçao and Bonaire.
23. van Buurt, G., & Debrot, A. O. (2011). Exotic and invasive terrestrial and freshwater animal species in the Dutch Caribbean (No. C001/12). IMARES.
24. Van Wilgen, N. J., Richardson, D. M., & Baard, E. H. (2008). Alien reptiles and amphibians in South Africa: towards a pragmatic management strategy. South African Journal of Science, 104(1-2), 13-20.

A)

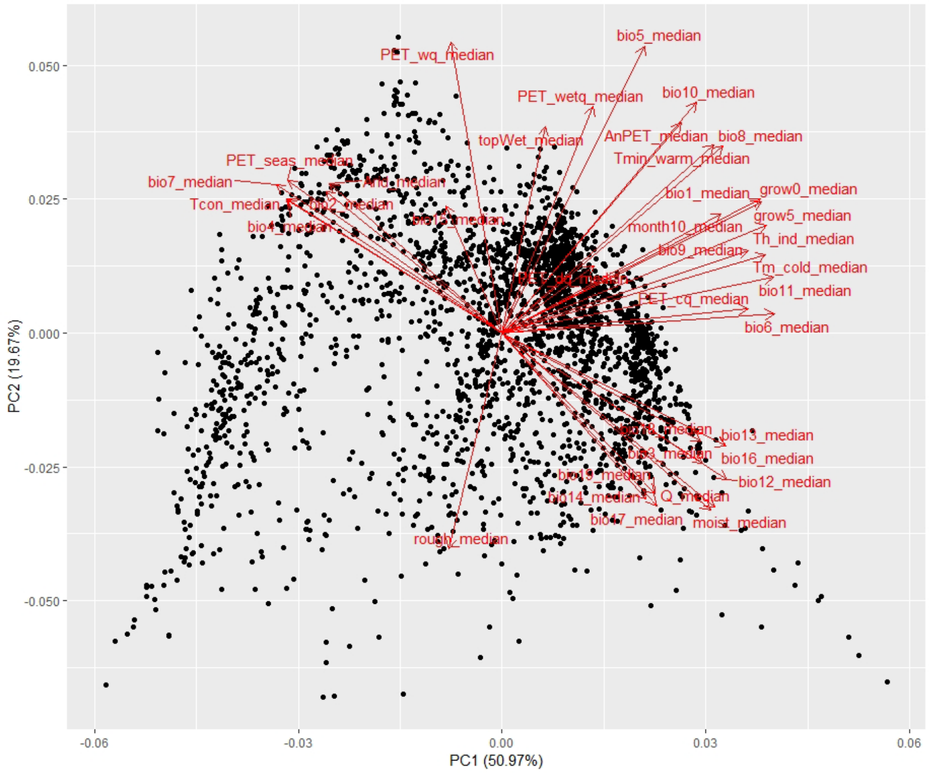

PC1 (50.97%)
PC2 (19.67%)
0.050
0.025
0.000
-0.025
-0.050
-0.06
-0.03
0.00
0.03
0.06
bio5_median
PET_wq_median
PET_wetq_median
bio10_median
topWet_median
AnPET_median
bio8_median
Tmin_warm_median
PET_seas_median
bio7_median
Arid_median
Tcon_median
bio2_median
bio4_median
bio1_median
grow0_median
grow5_median
month10_median
Th_ind_median
bio9_median
Tm_cold_median
bio11_median
PET_cq_median
bio6_median
bio13_median
bio16_median
bio12_median
Q_median
moist_median
bio17_median
bio14_median
rough_median


B)

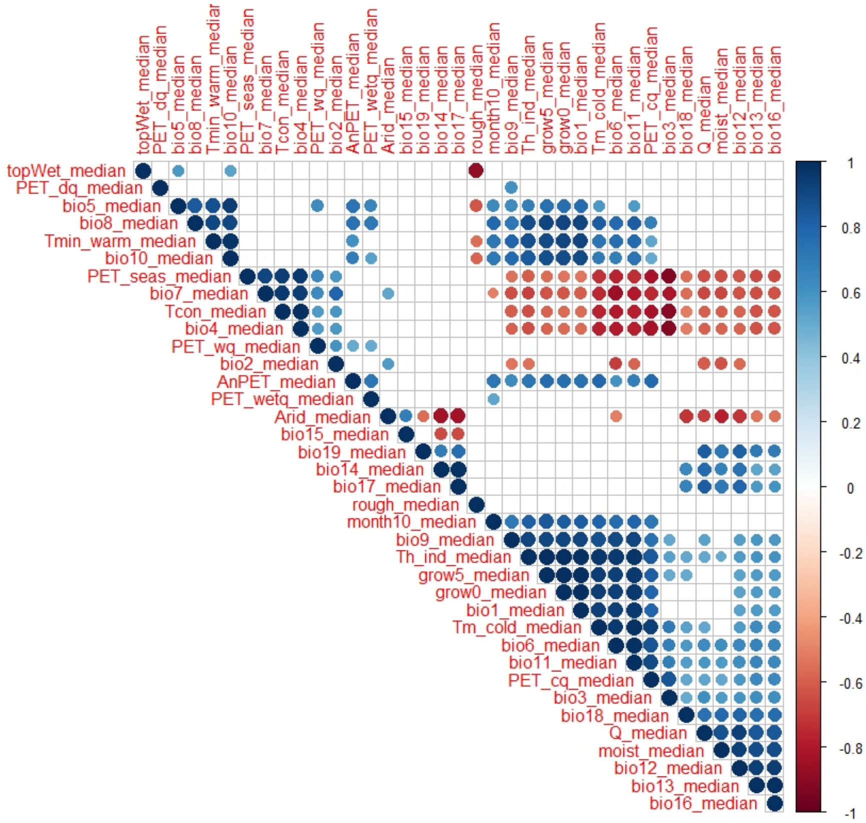

topWet_median
PET_dq_median
bio5_median
bio8_median
Tmin_warm_median
bio10_median
PET_seas_median
bio7_median
Tcon_median
bio4_median
PET_wq_median
bio2_median
AnPET_median
PET_wetq_median
Arid_median
bio15_median
bio19_median
bio14_median
bio17_median
rough_median
month10_median
bio9_median
Th_ind_median
grow5_median
grow0_median
bio1_median
Tm_cold_median
bio6_median
bio11_median
PET_cq_median
bio3_median
bio18_median
Q_median
moist_median
bio12_median
bio13_median
bio16_median
1
0.8
0.6
0.4
0.2
0
-0.2
-0.4
-0.6
-0.8
-1


1

**Figure S1**. Biplot and correlation matrix including all 37 climatic variables from Woldclim (Fick et al., 2017) and ENVIREM (Title & Bemmels, 2018) databases, which were included a scaled Principal Component Analysis (PCA). From this PCA, two environmental PC axes were derived and included in further analysis. A) Biplot of axis 1 (PC1) and axis 2 (PC2) of a PCAthat shows the variation among the median values of 37 climatic variables in 3061 species ranges. 1 point is 1 species range. PC1 aligns with variables indicating the level of seasonality within a species range (e.g., temperature seasonality; continentality and temperature range, negatively correlated to temperature in the coldest month). PC2 aligns with variables associated with elevational differences (e.g., terrain roughness index; SAGA-GIS topographic wetness index). We inverted both axes compared to the rotation depicted here. B) Correlation matrix of 37 WorldClim and ENVIREM variables for which the median per species range has been calculated. The variables are ordered to maximize congruence. Correlation coefficients > -0.5 and < 0.5 were excluded from this figure for clarity. Dot size indicates the strength of the correlations; the larger the dot, the closer the correlation coefficient is to 1 or –1. Blue dots indicate a positive correlation and red dots indicate a negative correlation among variables, and thus a parallel or opposite direction of the arrows in figure S1A, respectively.

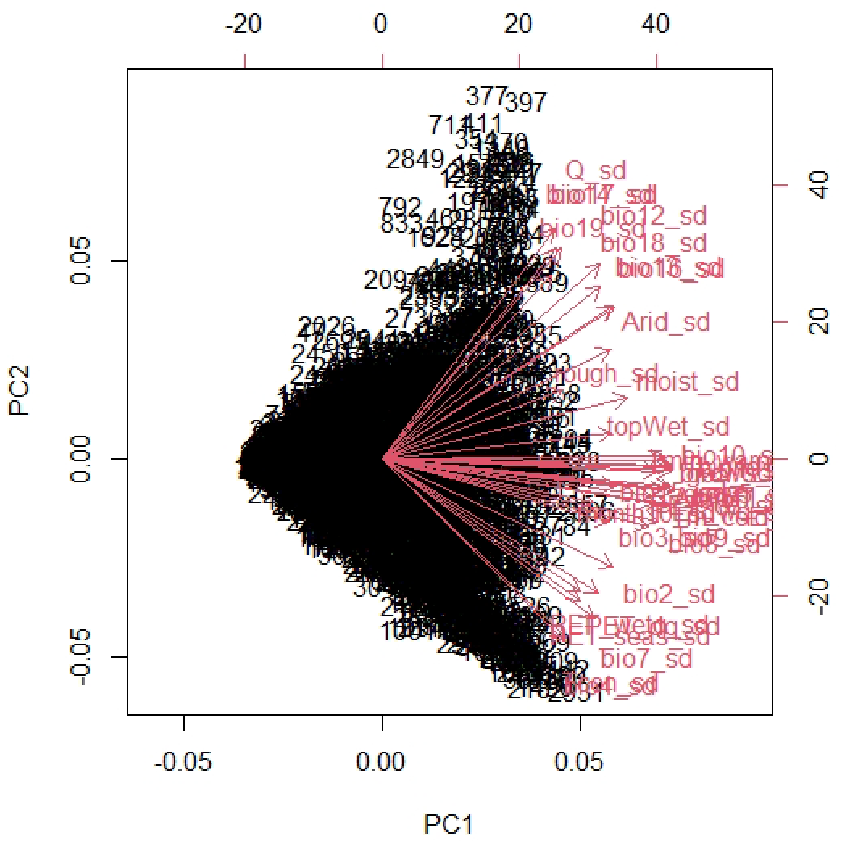


**Figure S2**. Biplot of WorldClim and ENVIREM climatic variables included in the climatic niche differentiation variable. The standard deviations of 37 climatic variables within 3056 species ranges were included in a principal component analysis (PCA). 1 point is 1 species range. PC1 aligns with all climatic variables, explaining a total of 63% of the overall variation. This means that if a standard deviation of for instance a temperature-based variable is high in a species range (i.e., a species experiences a variety of temperatures), also the standard deviations of other climatic variables increase in the same direction. And when the climatic standard deviations within a species range are high, the PC1 value for that species is also high. As it is expected that larger species ranges have more variable climates, PC1 was regressed against range size, taking the residuals of that relationship ($r = 0.65$, $t_{3054} = 47.45$, $P < 0.001$) as value for disproportional climatic niche differentiation/uniformity.

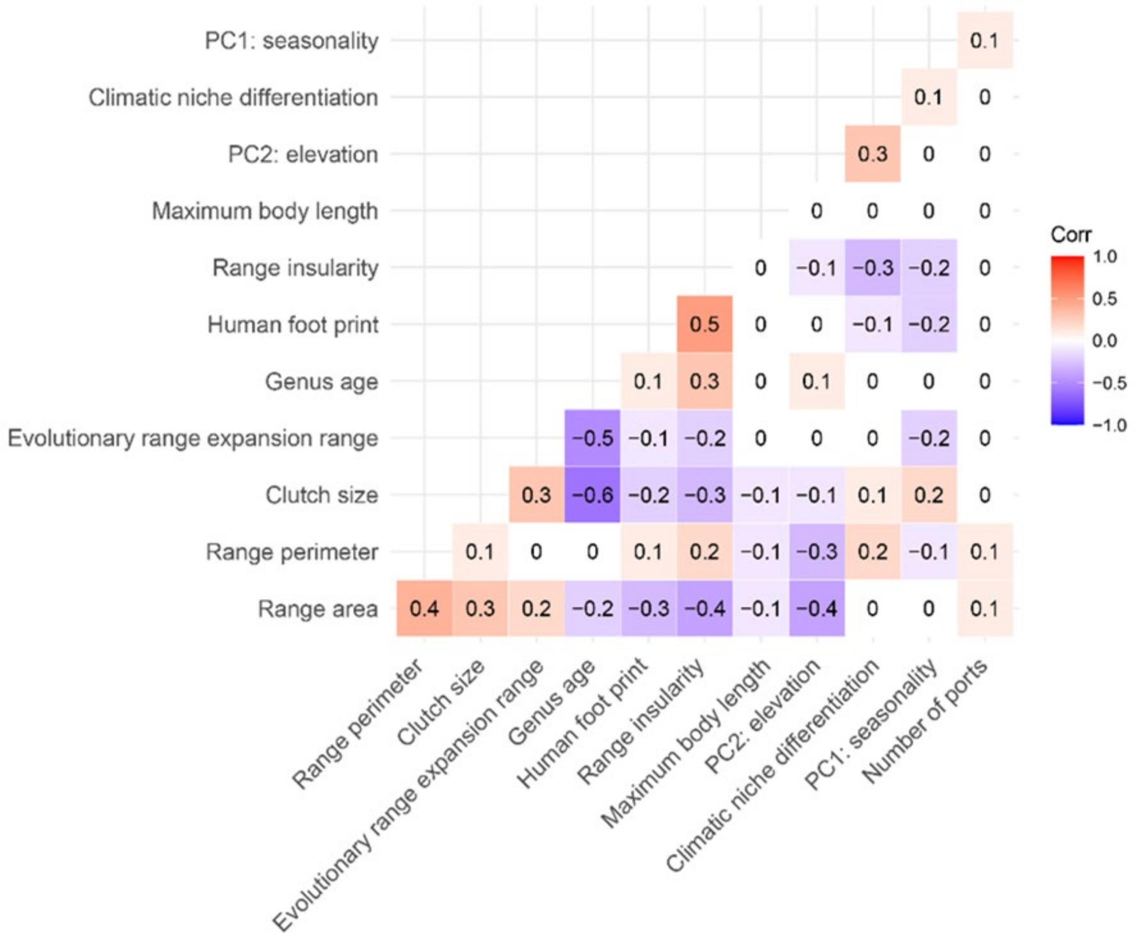


**Figure S3**. Correlations among predictor variables that were included in phylogenetic generalized linear models. Correlation matrix including all predictor variables. The brighter the color, the stronger the correlation between variables (i.e., closer to 1 or –1). Blue blocks indicate a negative relationship and red blocks indicate a positive relationship. The relatively low correlation coefficients among predictor variables indicate that there are no issues with collinear relationships that bias our model outputs (also indicated by a relatively low Variance Inflation Factor of <3, see main text)

**Table S2** Phylogenetic signal of predictor variables. Phylogenetic signal of discrete and continuous predictor variables. Phylogenetic signal is defined as Lamda (λ), ranging between 0 and 1, the higher the value, the stronger the signal ("fitContinuous" and "fitDiscrete" functions in *geiger* R package). We expected phylogenetic signal for some variables: Clutch size, maximum body length, diet type, and reproductive mode were phylogenetically imputed (see Rapacciuolo et al. 2019 for details). Furthermore, two variables were genus-specific variables (stem age, evolutionary range expansion range), hence inherently phylogenetically clustered.

| **Variable type** | **Predictor variable** | **Phylogenetic signal (λ)** |
|---|---|---|
| **Geographic** | **Range size** | 0.71816 |
| **Geographic** | **Range shape** | 0.59037 |
| **Geographic** | **Insularity** | 0.90954 |
| **Ecological** | **PC1: seasonality** | 0.81980 |
| **Ecological** | **PC2: elevation** | 0.70111 |
| **Ecological** | **Climatic niche differentiation** | 0.59214 |
| **Ecological** | **Diet type** | 0.94790 |
| **Ecological** | **Maximum body length** | 0.51557 |
| **Ecological** | **Clutch size** | 0.94050 |
| **Ecological** | **Reproductive mode** | 0.98400 |
| **Evolutionary** | **Genus age** | 1.00000 |
| **Evolutionary** | **Evolutionary range expansion rate** | 1.00000 |
| **Anthropogenic** | **Human footprint** | 0.72552 |
| **Anthropogenic** | **Number of ports** | 0.00007 |
